# HSV-1 orchestrates host RAP80 ubiquitination by ICP0 and UL36USP to promote viral survival

**DOI:** 10.1101/2025.06.10.658793

**Authors:** Jie Wang, Yujing Deng, Dongyue Zhu, Yi Chi, Fan Liu, Chunfu Zheng, Dong Xing, Xiaofeng Zheng

**Author notes:** To whom correspondence should be addressed &. These authors contribute equally to this work (Jie Wang, Yujing Deng).

## Abstract

Herpes simplex virus 1 (HSV-1) is a widespread human pathogen that establishes lifelong infections, ubiquitination is crucial for HSV - 1 to interact with the host and regulate its life activities. But the mechanisms of how HSV-1 dynamically modulates host proteins at different infection stages to promote its own survival remains challenging. Here we identified DNA damage response protein RAP80 as a critical target of HSV-1 ubiquitination enzymes ICP0 and UL36USP throughout HSV-1’s infection cycle. In early stages, RAP80 inhibits HSV-1 transcription by directly binding to the viral genome, blocking viral protein ICP4 binding to transcription factors. This interaction is regulated by phase separation via an intricate interplay between RAP80 and ICP0/ICP4. As infection progresses, the viral E3 ligase ICP0 degrades RAP80, dissolving phase separation, and allowing the formation of a mature viral replication compartment. In late infection stages, RAP80 is deubiquitinated by the viral deubiquitinase UL36USP, which restores cellular homeostasis and promotes HSV-1 survival. This process involves modulating the R-loop-cGAS-apoptosis pathway. These findings underscore the dynamic interplay between viral and host factors and the complex mechanisms used by HSV-1 to subvert host defenses, offering practical implications for the future development of antiviral strategies.

## Introduction

Herpes simplex virus 1 (HSV-1) affects 85% of adults, with lifelong, incurable infections. HSV-1 has a lifecycle consisting of immediate early, early, and late phases^1^. It hijacks host factors for transcription and replication purposes, including DNA damage response (DDR) proteins^2–4^. DDR pathways are essential for HSV-1 survival, but also inhibit viral replication and transcription^5–7^. Yet detailed mechanisms underpinning DDR’s inhibitory effects are unclear.

Following host cell entry, HSV-1 triggers innate immune responses. Innate immunity and DDRs are interconnected, with cells creating physiological links between the two. However, the precise mechanisms of how HSV-1 triggers such coordinated responses and antiviral effects are still being investigated. Innate immune and DDR pathways both induce apoptosis^8,9^, which is a double-edged sword in viral dynamics. Viruses may use apoptosis for viral particle release^10^, but also face the risk of being eliminated by excessive cell death^11,12^, which in turn protects neighboring cells^13^. This creates a paradox for viruses as they need to exploit cells for replication while evading host defenses. This has led to a coevolution between viruses and host cells^14^. Critically, understanding this complex viral-host equilibrium could facilitate new therapies.

HSV-1 exploits ubiquitination processes to modulate host factors to facilitate its life cycle. For instance, ICP0 (HSV-1 E3 ligase) degrades antiviral proteins like IFI16^15^, DNA-PK^16^, MORC3^17^, and RNF168^18^, raising concerns about over-apoptosis and its impact on cellular homeostasis, which may be essential for viral survival. However, ongoing research is predominantly focused on ICP0 substrates, and a comprehensive understanding of the cascades following host protein degradation and the resulting host-virus balance remains a challenge.

RAP80 is essential in the DDR pathway where it uses its two ubiquitin-interacting motif (UIM) to bind ubiquitin chains on H2A to help recruit the BRCA1-A complex to DNA damage sites, thus initiating repair^19–21^. However, RAP80 actions in balancing host-virus interactions and defenses remain poorly understood.

In this study, we highlight a crucial role for RAP80 as an unconventional participant in the intricate dance underlying viral-host interactions. To progress its life cycle, HSV-1 uses diverse ubiquitination strategies to regulate RAP80, thereby conferring it with several roles throughout the infection process. This sophisticated interplay highlights the dynamic and intricate nature of host-virus relationships and emphasizes the mechanisms that this virus uses to manipulate host factors for its own advantage.

### RAP80 suppresses HSV-1 transcription in early infection phases

We observed that RAP80 expression levels vary across distinct time points following HSV-1 infection (Fig. 1a). To investigate RAP80 roles within the HSV-1 life cycle, we first examined interactions between RAP80 and HSV-1 proteins. At 3 h post infection, IP-MS revealed that RAP80 interacted with several HSV-1 proteins, particularly the immediate-early proteins ICP0 and ICP4 (Extended Data Fig. 1a,b). Furthermore, a significant RAP80 colocalization with ICP0, ICP4, and ICP8 was observed (Fig. 1b). To investigate the impact of RAP80 on the HSV-1 life cycle, we performed Gene Ontology (GO) analysis on MS data. We observed RAP80 enrichment in viral transcription and gene regulation pathways (Extended Data Fig. 1c), thereby hinting at a role in HSV-1 transcription. HSV-1 gene transcription, including *TK and VP16* increased after *RAP80* knockdown (KD) (Fig. 1c and Extended Data Fig.1d), indicating that RAP80 inhibits HSV-1 transcription.

**Fig. 1.**
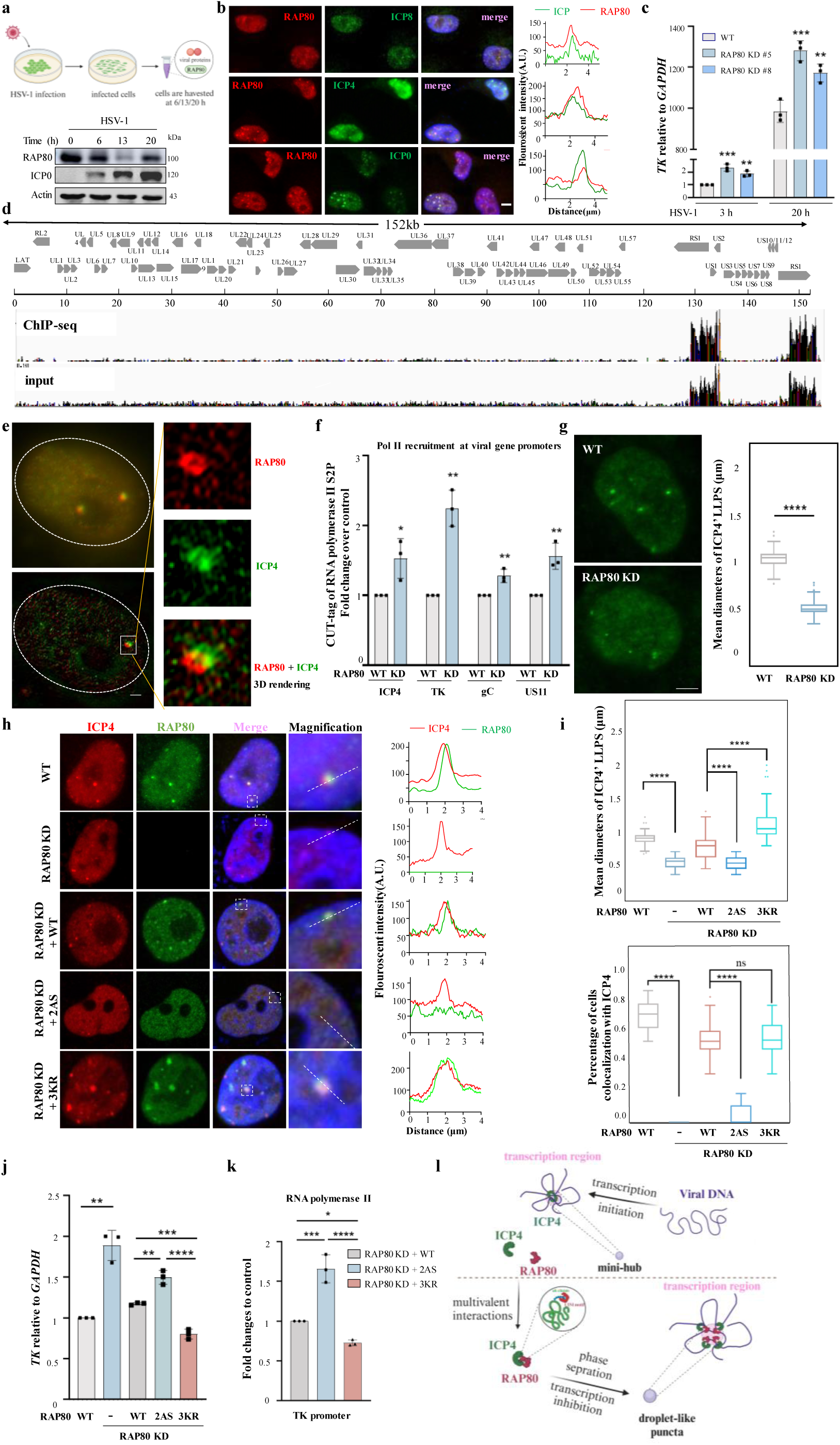
RAP80 suppresses HSV-1 transcription by orchestrating phase separation, which is driven by multivalent interactions during early HSV-1 infection stages. **a.** Endogenous RAP80 expression profiles across various post-HSV-1 infection stages were evaluated in HeLa cells exposed to HSV-1 at a multiplicity of infection (MOI) = 1. Cells were harvested at different time points post-infection, and RAP80 and ICP0 protein levels examined by immunoblotting with specific antibodies. **b.** Representative immunofluorescence images showing cellular RAP80 colocalization with the viral proteins ICP0, ICP4, and ICP8 in HeLa cells. Images were captured at 3 h post-infection with HSV-1 (MOI=1). Quantified RAP80 (red) and viral protein (green) fluorescence intensity values at colocalization sites were determined using ImageJ software. **c.** *RAP80* knockdown (KD) effects on viral *TK* expression were assessed in HeLa cells with depleted *RAP80* at 3 h or 20 h post-infection with HSV-1 (MOI=1). **d.** Specific RAP80 binding sites in the HSV-1 genome were investigated by ChIP-seq using ChIP DNA extracted from HeLa cells with a RAP80 antibody at 3 h post HSV-1 infection (MOI=1). Specific HSV-1 genomic regions, where ChIP-sequencing peaks aligned, are shown in a comprehensive plot. **e.** 3D-SIM was used to visualize RAP80 colocalization (red) with ICP4 (green) in HeLa cells infected with HSV-1 (MOI=1) for 3 h. **f.** *RAP80* KD effects on RNA polymerase II recruitment to viral gene promoters were evaluated in HeLa cells by ChIP assays using the RNA polymerase II S2P antibody. **g.** Representative immunofluorescence images showing ICP4 in WT and *RAP80* KD HeLa cells following infection with HSV-1 (MOI=1) for 3 h (left). ICP4 droplet diameters were measured in Imariz software (right). The graph represents data from three independent experiments and counts represent 50–100 infected cells. **h, i**. Immunofluorescence images showing *RAP80*-depleted HeLa cells re-expressing WT RAP80 or its mutant (*2AS* or *3KR*), which were infected with HSV-1 (MOI=1) for 3 h. Cells were stained with DAPI and ICP4 and RAP80-specific antibodies. (h) ICP4 droplet diameters and colocalization between different RAP80 and ICP4 forms were measured (i). **j.** RAP80 effects on *TK* expression were evaluated in *RAP80*-depleted HeLa cells plus reintroduced WT RAP80 or mutants (*2AS* or *3KR*). Cells were infected with HSV-1 (MOI=1) for 3 h and qRT-PCR assays performed to examine *TK* expression. **k.** RNA polymerase II recruitment to viral gene promoters was assessed in HeLa cells expressing WT RAP80 or mutants (*2AS* or *3KR*) in ChIP assays using a specific RNA polymerase II S2P antibody. **l.** The model shows multivalent interactions between RAP80 and ICP4 in phase separation formation (viral pre-replication foci). After HSV-1 infection, RAP80 binds to the viral transcription factor ICP4 and the viral genome through its UIM domain, driving phase separation formation. Data in panels **c, f, g, i-k** are presented as the mean ± standard error of the mean from three independent experiments. *P < 0.05, **P < 0.01 and ***P < 0.001. Data analysis was conducted using ordinary one-way and two-way analysis of variance (ANOVA) and Dunnett’s multiple comparisons tests. Exact P values are provided in source data. Scale bars, 5 μm.

### RAP80 suppresses HSV-1 transcription via phase separation

To elucidate how RAP80 inhibits HSV-1 transcription during early infection stages, we used ChIP-seq and detected that RAP80 directly bound to the viral genome, particularly where ICP4 bound (Fig. 1d and Extended Data Fig.1e). RAP80 binding sites are predominantly clustered in the promoter regions of ICP4-targeted genes (Extended Data Fig. 1e). A detailed IF results showed partial RAP80 and viral protein colocalization with spatial displacement (Fig. 1b). 3D-SIM showed that RAP80 and ICP4 form ring-like structures (Fig. 1e), suggesting these proteins have an antagonistic spatial distribution. Furthermore, CUT-tag assays revealed that *RAP80* KD increased RNA polymerase II binding at ICP4-target gene promoters, such as *TK, gC,* and *US11* (Fig. 1f and Extended Data Fig. 1f), suggesting RAP80 inhibits transcription by competing for ICP4 binding sites.

Given RAP80 colocalization with ICP4 in liquid droplet-like structures (foci, Fig. 1b), we investigated whether these foci represent phase separation. Using FRAP, we observed high RAP80 fluidity in cells (Extended Data Fig. 2a). High salt and 1,6-hexanediol levels reduced liquid-like RAP80 foci (Extended Data Fig. 2b). Real-time imaging also revealed RAP80 foci mobility and fusion/fission (Extended Data Fig. 2c,d). Also, *in vitro* phase separation assays confirmed the ability of RAP80 to form liquid droplets, which are disrupted by 1,6-hexanediol (Extended Data Fig.2e). These assays collectively indicate that RAP80 undergoes phase separation upon viral invasion. We further investigated the impact of RAP80 on ICP4’s phase separation. *RAP80* depletion significantly reduced ICP4 droplet sizes (Fig. 1g). To explored whether phase separation formation is regulated by interactions between RAP80 and ICP4, we conducted co-IP assays to examine RAP80-ICP4 interactions using *RAP80* truncation mutants and confirmed that the RAP80 UIM domain is key for binding ICP4 (Extended Data Fig.1g). The RAP80 UIM domain contains two conserved lysine residues (A88/112) which are crucial for ubiquitin binding (Extended Data Fig. 1h). We mutated these lysine to serine (termed *2AS*) residues and observed reduced binding to ICP4 (Extended Data Fig.1i). Additionally, we created stable cell lines using *RAP80 KD* cells expressing the *2AS* mutation, smaller ICP4 phase-separated foci were observed in cells stably expressing the *2AS* mutant, similar to cells with depleted *RAP80* (Fig. 1h, columns 2 and 4; Fig. 1i). These results underscore the indispensable role of the RAP80 UIM domain in facilitating phase separation.

Next, we investigated how RAP80 phase separation impacts its capacity to inhibit ICP4 transcriptional activity. The *RAP80 2AS* mutant had weaker effects on inhibiting ICP4 transcription when compared with the WT (Fig. 1j). Consistently, the binding affinity of *2AS* mutant to ICP4 target genes decreased (Extended Data Fig. 1j). Furthermore, enhanced RNA polymerase II binding to loci of these target genes in the *2AS* mutant was also observed (Fig. 1k). These results collectively indicate that RAP80 UIM-mediated phase separation is pivotal for its role in transcriptional repression.

These findings indicate that in early HSV-1 infection phases, the interaction between RAP80’s UIM domain and ICP4 is crucial for initiating phase separation. This interaction drive phase separation, which has an indispensable role in transcription inhibition of RAP80 (Fig. 1l).

### ICP0 ubiquitinates RAP80 to counteract RAP80’s antiviral effects

Notably, as infection progresses to mid-replication stages (6 h post-infection), we observed that the formation of a normal replication compartment (Fig. 2a, first panel), suggesting that dissolution of the RAP80-mediated phase separation is essential for the maturation of larger replication centers. ICP0 is known to counteract antiviral responses by ubiquitinating and degrading host antiviral proteins^18^. Here we identified significantly increased RAP80 ubiquitination levels after HSV-1 infection (Extended Data Fig. 3a). Moreover, we also infected cells with either WT HSV-1 or a variant lacking ICP0 ubiquitin E3 ligase activity (*ΔICP0-HSV-1*), and found that during the mid-replication stage (6 h), but not the late stage (18 h), cells infected with WT HSV-1 displayed a significant increase in K48- and K63-linked RAP80 ubiquitination (Extended Data Fig.3b). Notably, in cells infected with *ΔICP0-HSV-1*, RAP80 and ICP4 remained in a phase-separated state, which hindered the formation of a mature replication compartment (Fig. 2a, second panel). Subsequently, we extended the investigation by infecting WT and *RAP80*-KD HeLa cells with either the WT or *ΔICP0-HSV-1*, and then measured *TK* (viral gene) transcription levels and viral DNA production during early (2–3 h) and mid-replication stages (6–8 h) post-infection. In the presence of *ΔICP0-HSV-1*, *RAP80* KD significantly boosted *TK* gene transcription (Fig. 2b) and viral viability (Extended Data Fig. 3c). Thus, ICP0 appears to neutralize the antiviral impact of RAP80 via ubiquitination.

**Fig. 2.**
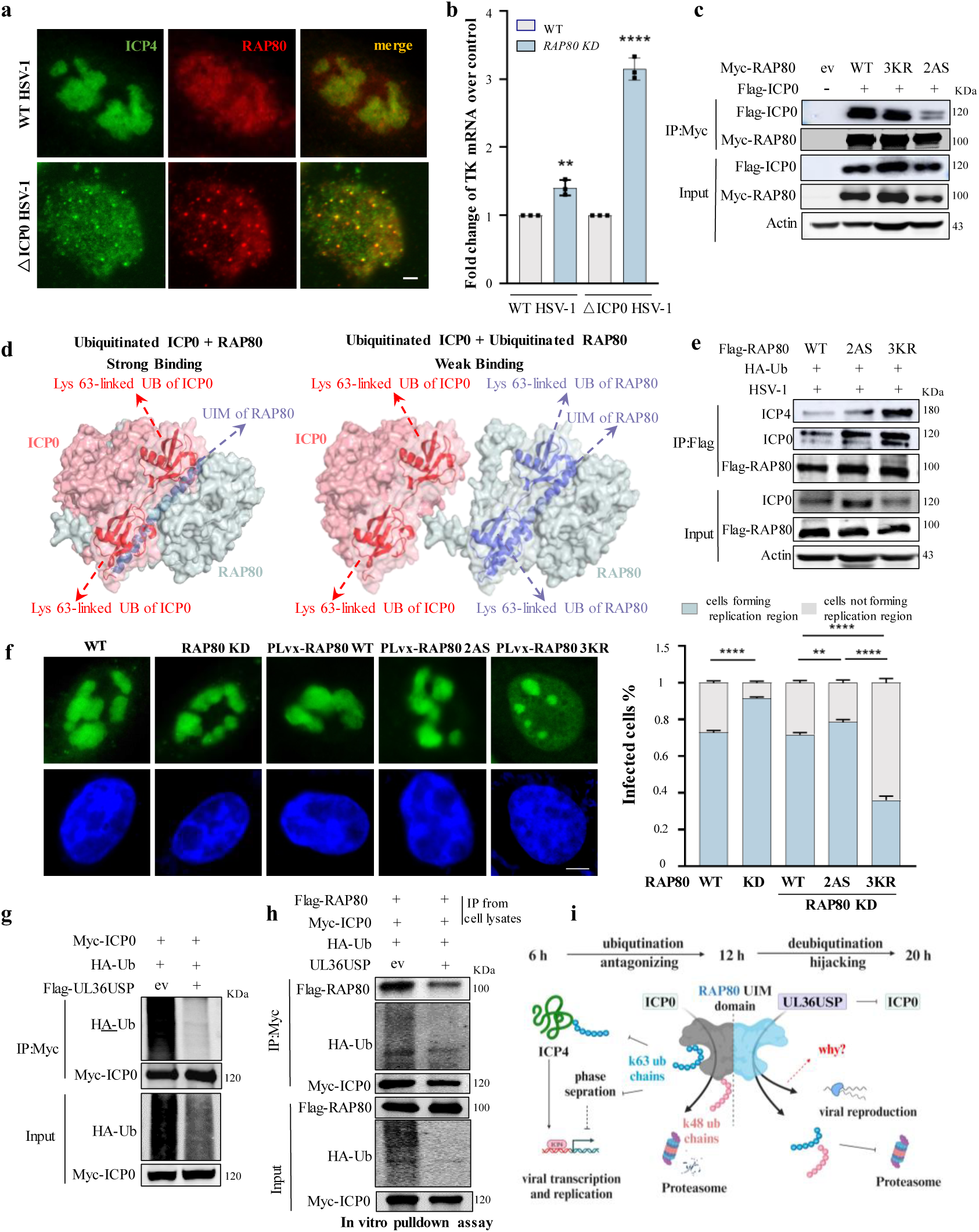
HSV-1 manipulates RAP80 function by targeting its ubiquitination via the viral proteins ICP0 and UL36USP at various infection stages. **a.** ICP0 enzyme effects on the establishment of the HSV-1 replication compartment were evaluated in HeLa cells by immunofluorescence microscopy. Cells were infected with either WT HSV-1 or an ICP0-deficient mutant strain (*ΔICP0 HSV-1*) (MOI=1) and incubated for 6 h. Subsequently, cells were stained with antibodies against ICP4 and RAP80. Representative immunofluorescence images are shown. **b.** ICP0 enzyme deficiency effects on *RAP80* depletion-promoted *TK* expression were explored in HeLa cells infected with either WT HSV-1 or *ΔICP0 HSV-1* (MOI=1) for 6 h. qRT-PCR was performed in triplicate to evaluate *TK* expression. **c.** Interactions between ICP0 and RAP80 mutants were examined using co-IP assays in HEK293T cells. Cells were co-transfected with Flag-tagged ICP0 and either Myc-tagged WT RAP80 or its mutants (*2AS* or *3KR*), and co-IP assays performed using indicated antibodies. **d.** Crystallographic analysis shows a significant binding pattern alteration in RAP80 following ubiquitination of its 3K sites (K75, 90, and 112). Post-ubiquitination, the RAP80 UIM domain binds to its own ubiquitination chains, attenuating its binding to ICP0. **e.** RAP80 ubiquitination effects on its association with viral proteins were examined in HEK293T cells by co-IP assays. Cells were co-transfected with Myc-RAP80 and either WT RAP80 or mutant variants (*2AS* and *3KR*) along with HA-tagged ubiquitin, and subsequently infected with HSV-1 for 6 h (MOI=1) before harvest. Cell lysates were subjected to IP with an anti-Flag antibody and analyzed by western blotting using anti-Flag, anti-ICP0, and anti-ICP4 antibodies. **f.** RAP80 and mutant effects on HSV-1 replication centers were observed by IF in HeLa cells infected with HSV-1 (MOI=1) for 6 h. **g.** The impact of UL36USP on ICP0-catalyzed ubiquitination was examined in HEK293T cells. Cells were co-transfected with Myc-tagged ICP0, HA-tagged ubiquitin, and Flag-tagged UL36USP, with cell lysates immunoprecipitated using an anti-Myc antibody, and analyzed by western blotting using anti-Myc and anti-HA antibodies. **h.** UL36USP effects on the interaction between RAP80 and ubiquitinated ICP0 were assessed in *in vitro* pull-down assays. Flag-tagged RAP80 immunoprecipitated from HEK293T cells was incubated *in vitro* with purified ubiquitinated ICP0 in the presence/absence of UL36USP protein, and then western blotting performed with indicated antibodies. **i.** The model shows the interplay of ubiquitination on RAP80 by ICP0 and UL36USP at different post-HSV-1 infection stages. During mid-stage, ICP0 catalyzes the K48- and K63-type ubiquitination of RAP80, resulting in RAP80 degradation and the dissolution of phase separation. In late infection stages, UL36USP deubiquitinates RAP80, thereby stabilizing RAP80 and facilitating viral propagation. Data in panels **b** and **f** are presented as the mean ± standard error of the mean from three independent experiments. **P < 0.01 and ***P < 0.001. Data analysis was conducted using ordinary one-way and two-way analysis of variance (ANOVA) and Dunnett’s multiple comparisons tests. Exact P values are provided in source data. Scale bars, 5 μm.

### ICP0 inhibits phase separation by disrupting RAP80-mediated multivalent interactions

We next investigated ICP0-promoted ubiquitination consequences on RAP80 biological roles. First, we infected cells with WT or *ΔICP0-HSV-1* and treated cells with MG132 (proteasome inhibitor, control). At 6 h post-infection, we observed RAP80 degradation via the ubiquitin-proteasome system in an ICP0-dependent manner (Extended Data Fig. 3d). Additionally, ICP0 promoted the K63-linked ubiquitination of RAP80 (Extended Data Fig.3b), leading us to investigate the functional implications of this modification. We identified three lysine residues (K75, K90, and K112) in the UIM domain of RAP80 and constructed a triple lysine mutant (*3KR*) to examine if ICP0 could still induce K63-linked ubiquitination at these sites and see how this would affect RAP80’s antiviral functions. ICP0 selectively promoted K63-linked, but not K48-linked ubiquitination at these residues (Extended Data Fig. 3e). Subsequently, we analyzed the impact of K63-linked ubiquitination on the interaction with ICP0. Given that interactions between ICP4 (Extended Data Fig.1i), ICP0 (Fig. 2c), and RAP80 depend on the RAP80 UIM domain, mutation of UIM’s conserved sites (A88/112) in the *2AS* mutant reduced its interactions with ICP0 and ICP4. Considering the location of K63-linked ubiquitination sites in the UIM domain, we hypothesized that RAP80 ubiquitination at K75, K90, and K112 sites enhances RAP80’s K63-linked ubiquitin chain binding to the UIM domain, potentially shielding it and inhibiting interactions with ICP0 and ICP4 (Fig. 2d). We tested this hypothesis using HSV-1-infected cells with WT and the *3KR RAP80* mutant and found that the mutant showed enhanced interactions with ICP0 and ICP4 when compared with WT RAP80 (Fig. 2e).

To further ascertain whether RAP80 selectively interacts with the ICP0 ubiquitin chain, mirroring its recognized role in DDR pathways where it binds to histone ubiquitin chains, we performed *in vitro* pull-down assays (Extended Data Fig.3f) and found that *2AS* mutant binding capacity to ICP0 was reduced, whereas no significant difference in binding capacity was observed between WT RAP80 and the *3KR* mutant when interacting with ICP0 (Extended Data Fig. 3g). Moreover, interactions between ubiquitinated RAP80 or its mutants and ubiquitinated ICP0 revealed that RAP80 *2AS* mutant interaction with ICP0 remained unchanged relative to WT, whereas the *3KR* mutant exhibited enhanced binding capacity to ICP0 (Extended Data Fig. 3h), supporting *in vivo* results (Fig. 2e). These findings suggest that RAP80 binds to the ICP0 ubiquitin chain, facilitating multivalent interactions. Thus, RAP80 ubiquitination by ICP0 masks its UIM domain, reducing interactions with ICP0 and ICP4.

Next, we stably expressed the *3KR RAP80* mutant in *RAP80*-depleted cells to investigate its physiological effects. In comparison to WT cells, the *3KR* mutant formed larger phase-separated clusters with ICP4 (Fig. 1h,i) and increased associations with the HSV-1 genome (Extended Data Fig. 1j), thereby reducing RNA polymerase II binding capacity to the HSV-1 genome (Fig. 1k) and enhancing HSV-1 transcription suppression (Fig. 1j). During the mid-stage HSV-1 life cycle, the *3KR* mutant notably reduced mature replication compartments (Fig. 2f), yet these centers retained a larger circular morphology when compared with typical phase-separated structural dimensions. We further examined RAP80 self-interaction at different viral infection stages. Early in HSV-1 infection (3 h post-infection), RAP80 exhibited strong self-interactions, correlating with additional phase separation (Fig. 1b). In terms of mid-stage HSV-1 infection (6 h post-infection), WT RAP80 and the *2AS* mutant displayed reduced self-interactions, while the *3KR* mutant sustained strong self-interactions (Extended Data Fig.3i), consistent with phase separation (Fig. 1h).

Taken together, ICP0 emerges as a key viral protein counteracting antiviral RAP80 effects, and accomplishing this role by orchestrating RAP80 ubiquitination.

### UL36USP restore RAP80 during late HSV-1 infection stages

To investigate the regulatory mechanisms underlying the restored RAP80 stability in late infection phases observed in Fig. 1a, we examined RAP80 ubiquitination status at different times post HSV-1 infection. When compared to the early phase (6 h post-infection), a marked reduction in K63- and K48-linked RAP80 ubiquitination was observed in the late phase (18 h post-infection) (Extended Data Fig.3b). To pinpoint factors responsible for RAP80 deubiquitination in late HSV-1 infection, we investigated UL36USP, a viral deubiquitinase active during this phase^22^. UL36USP deubiquitinated RAP80 (Extended Data Fig. 4a,b) and interacted with the protein (Extended Data Fig.4e). A catalytic site mutation (*UL36C40A*) in UL36USP eliminated its deubiquitination ability (Extended Data Fig.4c) and attenuated the interaction with RAP80 (Extended Data Fig.4f). Notably, UL36USP removed K48- and K63-linked ubiquitination from RAP80 (Extended Data Fig.4d), consistent with physiological conditions (Extended Data Fig. 3b). These findings collectively underscore the pivotal role of UL36USP in restoring RAP80 stability via deubiquitination during late HSV-1 infection stages.

### UL36USP antagonizes ICP0-mediated RAP80 ubiquitination

To understand the interplay between ICP0 and UL36USP in regulating RAP80 ubiquitination during late HSV-1 infection stages, we observed that UL36USP overexpression inhibited ICP0-induced RAP80 ubiquitination (Extended Data Fig.4g). Notably, under physiological conditions, RAP80 ubiquitination levels increased during mid-infection but declined at late stages (Extended Data Fig.4h), suggesting that when ICP0 and UL36USP are present, UL36USP deubiquitination effects predominate. Furthermore, during late HSV-1 infection stages, UL36USP significantly reduced the interaction between ICP0/4 and RAP80 (Extended Data Fig.4h), suggesting that UL36USP deubiquitinates RAP80 and inhibits its binding to ICP0/4. To elucidate this regulatory mechanism, we observed that UL36USP also interacted with the RAP80 UIM domain (Extended Data Fig.4i), overlapping with ICP0/4’s binding site. UL36USP’s deubiquitination likely strengthens this interaction by re-exposing the UIM domain. We also observed that UL36USP reduced the ICP0–RAP80 interaction by binding to the RAP80 UIM domain and deubiquitinating ICP0 (Fig. 2g,h). This dual action enhances the interaction between UL36USP and RAP80, thereby emphasizing UL36USP’s pivotal role in stabilizing RAP80 through deubiquitination.

To investigate the influence of UL36USP-mediated deubiquitination on RAP80 function, we examined UL36USP effects on RAP80’s phase separation, damage site recruitment, and viral genome binding. Initially, exogenous (Extended Data Fig.5a) and endogenous stable UL36USP overexpression (Extended Data Fig.5b) promoted RAP80 phase separation, aligning with increased RAP80 self-interactions (Extended Data Fig.5c) and indicating that UL36USP’s deubiquitination of RAP80’s UIM domain strengthens its protein interaction capacity. RAP80 capacity to associate with the viral genome was also restored during late stages (Extended Data Fig.5e). Therefore, RAP80 is revitalized via UL36USP actions during late HSV-1 infection stages (Fig. 2i).

### RAP80 suppresses HSV-1-induced apoptosis to enhance HSV-1 survival

We observed that *RAP80* KD reduced HSV-1 titers (Fig. 3a). To elucidate the physiological significance of UL36USP-mediated deubiquitination of RAP80 in HSV-1 infections, we used GFP-tagged HSV-1 in a 48-h infection period, allowing for multiple viral replication and lysis cycles, to detect viral fluorescence intensity. Importantly, the *RAP80* KD reduced HSV-1 fluorescence (Fig. 3b), indicative of decreased viral replication. These insights suggest that RAP80’s timely restoration at late stages is equally crucial for HSV-1 survival.

**Fig. 3.**
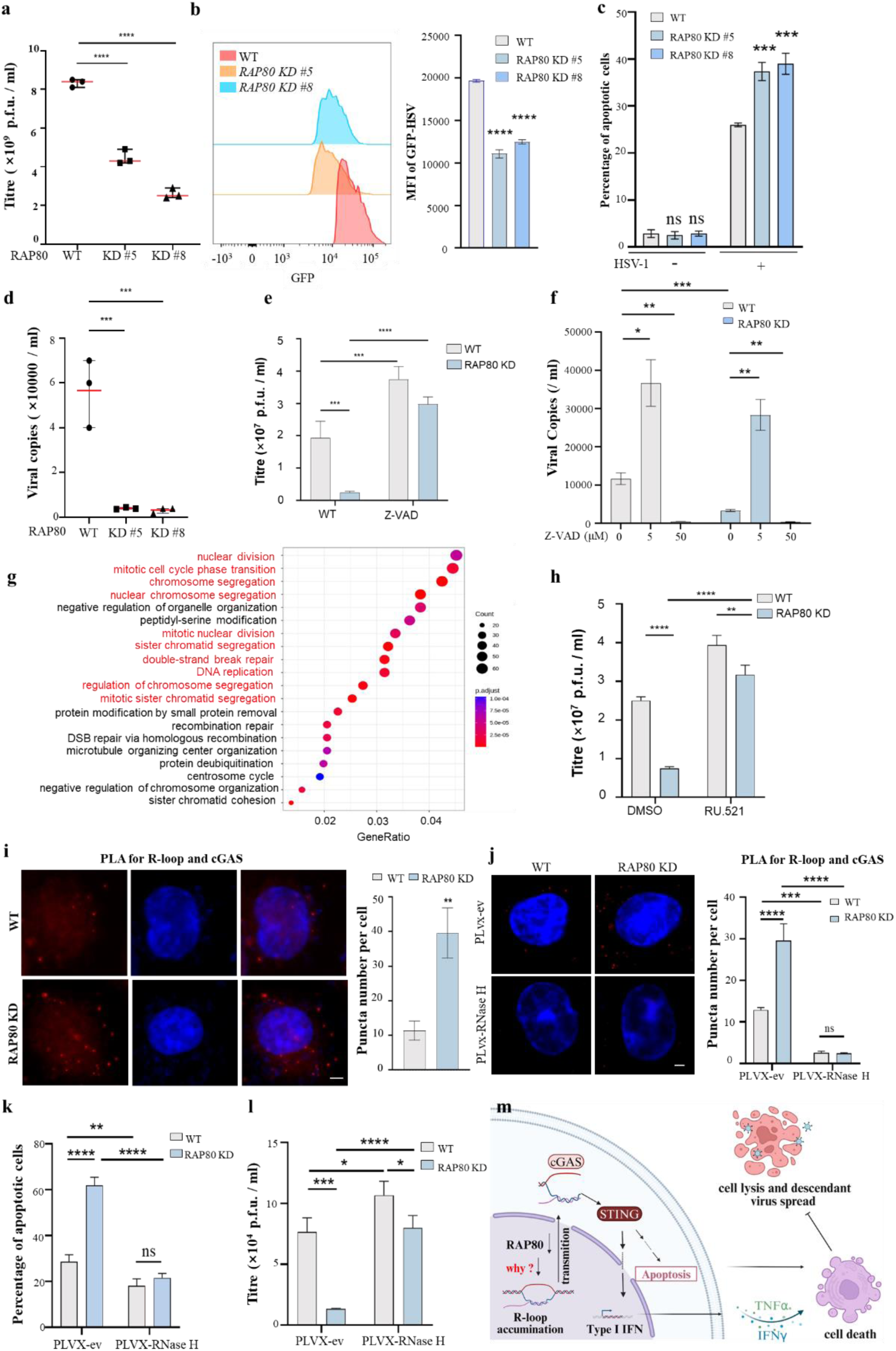
RAP80 mitigates late stage HSV-1 replication inhibition by disrupting the R-loop-cGAS-apoptosis-axis. **a.** *RAP80* depletion effects on HSV-1 replication during late infection stages were monitored using plaque assays in HeLa cells with *RAP80* knockdown (KD) at 20 h post HSV-1 infection (MOI=1). WT HeLa cells under identical conditions served as controls. **b.** WT HeLa and *RAP80* KD cells were infected with HSV-1-GFP for 20 h, after which, viral GFP mean fluorescence intensity (MFI) was quantified using flow cytometry to determine *RAP80* depletion effects on HSV-1 amplification. **c.** WT and *RAP80* KD HeLa cells were infected with HSV-1 (MOI=1) for 20 h, after which RAP80 effects on apoptosis were evaluated by Annexin V and propidium iodide (PI) staining. WT HeLa cells under identical condition served as controls. **d.** *RAP80* KD effects on viral replication were measured by infecting WT and *RAP80* KD HeLa cells with HSV-1 (MOI=1). At 20 h post-infection, supernatants were harvested and incubated with BHK cells and finally subjected to plaque assays to measure HSV-1 replication. **e.** The impact of the apoptosis inhibitor Z-VAD on HSV-1 amplification in *RAP80* KD HeLa cells was examined. WT and *RAP80* KD cells were pretreated with 5 μM Z-VAD or control conditions before infection with HSV-1. Viral replication was quantified using plaque assays. **f.** WT and *RAP80* KD HeLa cells were exposed to a Z-VAD concentration gradient (5–50 μM), after which plaque assays were conducted to assess varying Z-VAD concentration effects on HSV-1 replication efficiency. **g.** Gene enrichment analysis was conducted using RNA sequencing data to identify differentially expressed genes in *RAP80* KD HeLa cells infected with HSV-1 at 20 h post-infection (MOI=1). **h.** WT and *RAP80* KD HeLa cells were treated with 5 μM of RU.521 (cGAS inhibitor) prior to infection with HSV-1. HSV-1 replication was then measured using plaque assays. **i.** *RAP80* KD effects on the interaction between cGAS and R-loops was assessed using PLA in WT and *RAP80* KD HeLa cells. After infection with HSV-1 (MOI=1) for 20 h, PLA was performed to investigate *in situ* interactions between cGAS and R-loops (using the S9.6 antibody). **j.** PLA was performed to investigate RNase H effects on *RAP80* depletion-promoted R-loops in HeLa cells. WT and *RAP80* KD HeLa cells with/without overexpressed RNase H were infected with HSV-1 (MOI=1) for 20 h, and PLA conducted to examine interactions between cGAS and R-loops (S9.6 antibody). **k, l.** WT and *RAP80* KD HeLa cells with/without overexpressed RNase H were infected with HSV-1 (MOI=1) for 20 h, after which flow cytometry was conducted to examine apoptosis levels (k), and plaque assays performed to quantify viral replication (l). **m.** The model shows disrupted cellular homeostasis by HSV-1 following *RAP80* KD. *RAP80* depletion leads to R-loop accumulation and transmission to the cytoplasm. In the cytoplasm, R-loops are recognized by cGAS, activating innate immunity pathways and inducing apoptosis, which attenuates the viral lytic cycle. Data are presented as the mean ± standard error of the mean from three independent experiments. *P < 0.05, **P < 0.01 and ***P < 0.001. Data analysis was conducted using ordinary one-way and two-way analysis of variance (ANOVA) and Dunnett’s multiple comparisons tests. Exact P values are provided in source data. Scale bars, 5 μm.

We next investigated how RAP80 regulates HSV-1 survival. At 20 h post-infection, *RAP80* KD induced higher apoptosis rates (Fig. 3c). Additionally, progeny virion release was drastically reduced (Fig. 3d) and increased danger signals (e.g., ATP) were observed in cells (Extended Data Fig.6a). To assess if RAP80 regulates HSV-1 survival by affecting apoptosis, we infected cells with GFP-tagged HSV-1 for 16 h, allowing for initial viral replication, but without cell lysis. We observed that the *RAP80* KD led to faster HSV-1 replication and higher apoptosis rates (Extended Data Fig. 6b). Addition of the apoptosis inhibitor Z-VAD reduced HSV-1-induced apoptosis in a dose-dependent manner, with higher doses (50 μM) more effective than lower doses (5 μM) (Extended Data Fig.6c). In low Z-VAD-treated cells, HSV-1 titers increased in WT and *RAP80*-depleted cells (Fig. 3e, Extended Data Fig.6d). To examine if RAP80 effects on viral titers were mediated by its impact on progeny virus release via apoptosis, we treated cells with high and low Z-VAD concentrations and quantified progeny viruses. As expected, only moderate apoptosis (low Z-VAD dose) allowed efficient virus release, while excessive apoptosis and its inhibition reduced titers (Fig. 3f).

### RAP80 suppresses R-loop-cGAS-triggered apoptosis

We used RNA-seq to explore how RAP80 regulates apoptosis in HSV-1-infected cells. *RAP80* KD increased gene expression linked to cell division, cell cycle, and chromosomal segregation pathways (Fig. 3g). We hypothesized that RAP80 affects centromere function, which is key for chromosomal segregation and genomic stability^23^. Indeed, cells lacking *RAP80*, post-HSV-1 infection, showed more genomic instability (Extended Data Fig.7a). Notably, the centromere-specific histone CENP-A^23^, which normally localizes to the nucleus, was detected in the cytoplasm where it colocalized with cGAS (Extended Data Fig.7b). This interaction triggered the activation of cytoplasmic innate immune pathways, leading to increased IFN-β and TNF-α expression (Extended Data Fig.7f). These results suggest that *RAP80* KD exacerbates genomic instability, which in turn, is sensed by cytoplasmic cGAS, thereby activating innate immune responses.

To determine whether impaired centromere function and genomic instability, caused by *RAP80* KD along with innate immune pathway activation, were responsible for apoptosis during HSV-1 infection, we used the cGAS-specific inhibitor RU.521. cGAS inhibition reduced apoptosis levels induced by *RAP80* KD (Extended Data Fig.6e), while concurrently, HSV-1 titers and cell survival rates increased significantly (Fig. 3h, Extended Data Fig.6f). These results underscore the pivotal role of cGAS in mediating apoptosis following *RAP80* KD.

To investigate how genomic instability activates cGAS and examine its impact on innate immunity and cellular homeostasis, we identified nuclear factors that trigger cGAS. No pronounced overlap between cytoplasmic double-stranded DNA (dsDNA) and cGAS (Extended Data Fig.7c,d). Instead, colocalized cGAS and R-loops were observed after HSV-1 infection (Extended Data Fig.7d), suggesting R-loops as potential initiators of cGAS activation. R-loops are RNA and DNA hybrids which are involved in DNA damage repair, transcription-coupled replication, and centromere function^24^. Therefore, centromere instability induced by *RAP80* KD likely led to R-loop accumulation and leakage into the cytoplasm, thereby affecting cGAS-mediated innate immune responses and apoptosis. To validate this hypothesis, we conducted DNA-RNA Immunoprecipitation (DRIP) assays and found that cGAS bound R-loops more effectively after *RAP80* depletion at 20 h post HSV-1 infection (Extended Data Fig.7e). Furthermore, *RAP80* KD heightened cGAS-R-loop interactions (Fig. 3i). To test if R-loops activate cGAS innate immune pathways, we stably overexpressed RNase H, an enzyme mediating R-loop cleavage. RNase H overexpression reversed R-loop buildup in *RAP80* KD cells (Fig. 3j). Reduced R-loop levels were associated with decreased innate immune pathway activation (Extended Data Fig.7g), reduced apoptosis rates (Fig. 3k), and increased HSV-1 titers (Fig. 3l). These collective findings highlight the pivotal role of the R-loop-cGAS axis in modulating innate immune responses, which in turn influences apoptosis and HSV-1 survival in the context of an *RAP80* KD (Fig. 3m).

### RAP80 maintains centromere stability by suppressing R-loop and RAD52-mediated DNA damage responses at centromeres

RAP80 depletion boosted associations between R-loops and cGAS, prompting us to explore how RAP80 regulated R-loop accumulation. RAP80 genomic distribution patterns showed that RAP80 was localized to centromeres throughout early and late stages (Extended Data Fig.8a). Additionally, DNA sequences bound by RAP80 were rich in α-satellite and centromere-specific elements (Fig. 4a, Extended Data Fig.8b), suggesting direct engagement with centromeric regions. Next, we investigated if RAP80 preserves centromere stability by regulating R-loop function. Upon *RAP80* depletion, more R-loops were observed at centromeres (Fig. 4b), suggesting that RAP80 prevents R-loop accumulation at centromeres.

**Fig. 4.**
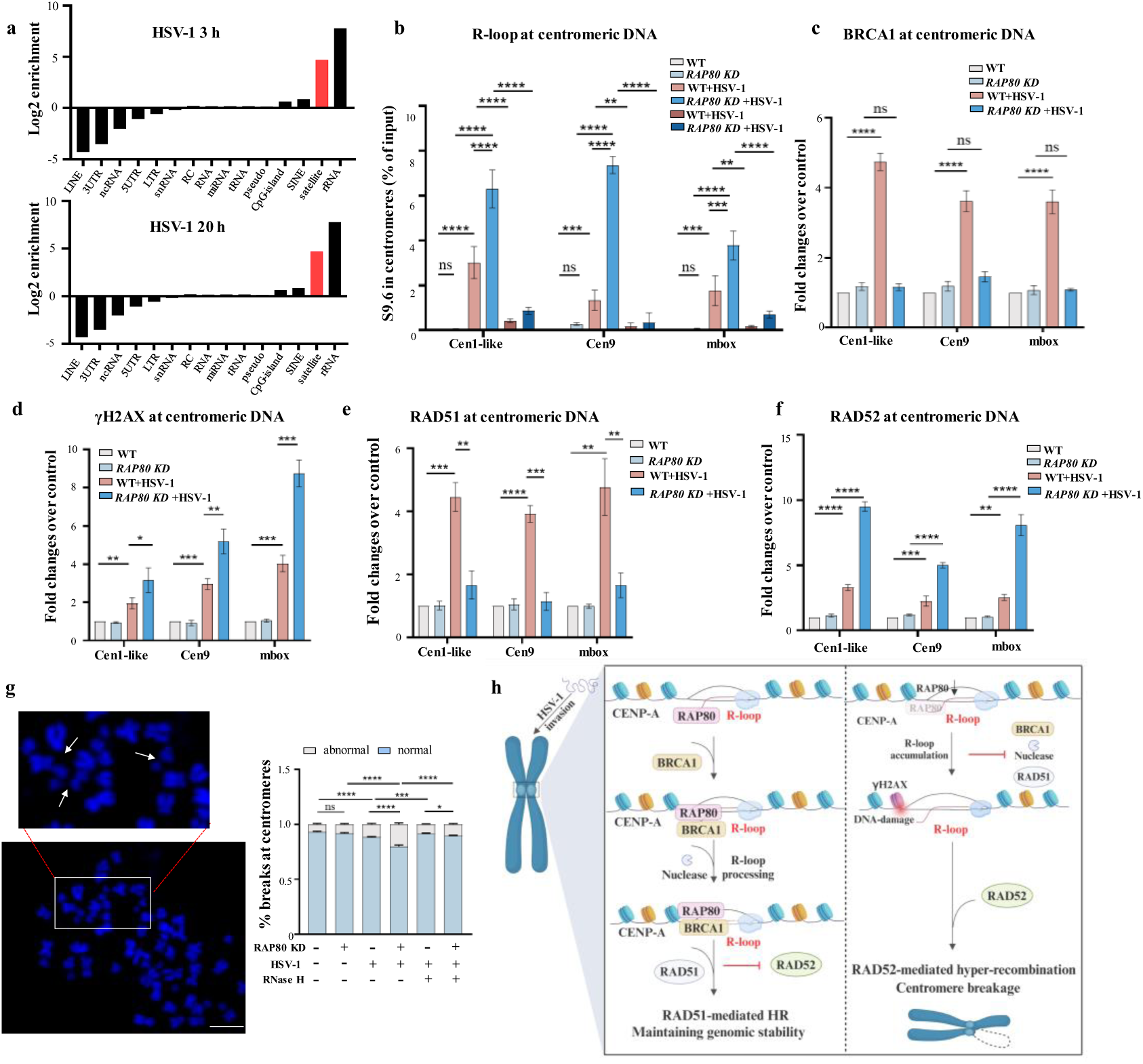
RAP80 functions as a guardian of centromere stability by suppressing DNA damage triggered by R-loop accumulation. **a.** ChIP-seq analysis was performed using an RAP80-specific antibody in HeLa cells, with ChIP DNA collected at 3 h and 20 h post-infection (MOI=1). Filtered peaks from ChIP-seq data were then annotated and analyzed for RAP80 binding patterns. **b.** R-loop accumulation at centromeres was evaluated in WT and *RAP80* KD HeLa cells infected with HSV-1 (MOI=1) for 20 h post-infection, in the presence/absence of RNase H, using CUT-tag– quantitative PCR (CUT-tag-qPCR) assays with an anti-S9.6 antibody to analyze the binding profiles of various centromeric DNA, Cen1-like, Cen9, and mbox. **c–j.** Protein recruitment, including BRCA1(c), γH2AX (d); RAD51 (e), RAD52 (f) to centromeres, was evaluated in WT and *RAP80* KD HeLa cells infected with HSV-1 (MOI=1) for 20 h. CUT-tag-qPCR assays were used to analyze protein binding profiles to different genes, including *Cen1-like, Cen9,* and *mbox*. **g**. Representative images showing chromosome spreads from HeLa cells with WT or depleted *RAP80* in RNase H overexpressing cells infected with HSV-1 20 h post infection (MOI=1). White arrows indicate centromeric breaks. **h.** The model highlights a pivotal role for RAP80 in maintaining centromere stability via degraded R-loops. RAP80 degrades R-loops by recruiting BRCA1 and nucleases. *RAP80* knockdown (KD) leads to R-loop accumulation and DNA damage at centromeres. This accumulation not only threatens centromere structural integrity but also disrupts finely tuned DNA damage repair processes. RAP80 plays a role in pathway selection for centromeric DSB repair. In its absence, DSBs are repaired by the RAD52-mediated pathway, and RAD51 is excluded from centromeric DSBs. The RAD52-mediated repair pathway results in uncontrolled homologous recombination, which is coupled to R-loop and DSB accumulation, and also prevents CENP-A deposition and promotes transcription. These combined effects lead to centromeric instability. Data in panels **b-g** are the mean ± standard error of the mean from three independent experiments. *P < 0.05, **P < 0.01 and ***P < 0.001. Ordinary one-way and two-way analysis of variance (ANOVA) and Dunnett’s multiple comparisons tests. Exact P values are provided in source data. Scale bars, 5 μm.

Since RAP80 lacks nuclease activity, we sought to elucidate how RAP80 facilitates R-loop degradation, which is primarily mediated by RNase H, XPG, and SETX nucleases^24^. Also, DDR factors such as BRCA1 regulate R-loop degradation at centromeres^25^. As RAP80 is vital for BRCA1-dependent DNA damage repair^19,20^, we investigated if R-loop inhibition by RAP80 was also mediated by BRCA1. *RAP80* depletion reduced BRCA1 binding to centromeres (Fig. 4c). Furthermore, R-loop formation was a prerequisite for BRCA1 recruitment (Extended Data Fig.8c), suggesting that RAP80 is a key factor for BRCA1 recruitment to centromeric R-loops. But RAP80 associated with centromeres without R-loop involvement (Extended Data Fig.8d). Collectively, these results demonstrate that RAP80 promotes R-loop degradation by recruiting BRCA1 and nucleases, thereby explaining why failure to restore RAP80 expression leads to HSV-1-induced R-loop accumulation and cGAS-innate immunity-apoptosis responses.

Given the link between centromeric R-loops and DNA damage^25,26^, *RAP80*-depleted cells contain more R-loops and increased γH2AX centromere binding after HSV-1-infected (Fig. 4d). Furthermore, R-loop degradation by RNase H significantly diminished γH2AX binding at centromeres (Extended Data Fig.8e), suggesting that R-loops induce DNA damage at centromeres. To explore how R-loop-induced DNA damage at centromeres is repaired, and how RAP80 acts in this process, we examined the centromeric binding of two crucial proteins in HR repair: RAD51 and RAD52. RAD51, not RAD52, mainly bound to centromeres (Fig. 4e,f). In contrast, without *RAP80*, RAD52 bound instead of RAD51 (Fig. 4e,f). And both RAD51 and RAD52 bound to centromeres in a manner that depends on the accumulation of R-loop (Extended Data Fig 8f,8g). RAD52 functions in single-strand annealing (SSA), thus excess of which leads to uncontrolled recombination, genomic instability, and even deleterious effects^27^. Given that this pathway causes genomic instability^25–27^, we also found that *RAP80* depleted-HSV-1-infected cells exhibited centromere dysfunction, with frequent chromosome breaks at centromeres (Fig. 4g). These findings indicate that R-loop accumulation and the RAD52-mediated repair pathway lead to centromere instability and subsequent breaks, RAP80 prevent all of above. (Fig. 4h).

### RAP80 modulates HSV-1 genome docking at host centromeres by regulating centromere stability and transcription status

After a virus invades host cells, virus-host genome interactions are mediated by both virus- and host-encoded proteins. Previous studies have shown that transcriptionally active euchromatin is more likely to serve as a viral docking site in the host genome^28^. Using high throughput chromosome conformation capture technology, we calculated HSV-1 preferences for centromeric regions. In early stages, fewer viral genome interactions were observed at centromeres (Extended Data Fig.9a, Fig. 5a), likely due to the heterochromatic nature of centromeres. However, between 9 h and 15 h post-invasion, the proportion of viral dockings at centromeres increased, followed by a decline between 15 h and 24 h (Fig. 5a,b). Interestingly, this pattern was inversely correlated with RAP80 protein levels, suggesting that RAP80 may regulate these interactions. However, interactions between HSV-1 and the host genome were not completely abolished after *RAP80* KD (Fig. 5c), indicating that RAP80 is not a direct mediator of these interactions, but may regulate this process via other pathways. Further analyses showed that after *RAP80* knockout, RNA polymerase II binding at centromeres, along with centromeric RNA transcription, increased dramatically (Fig. 5d,e). This shift from a transcriptionally inactive heterochromatic state to a more active euchromatic state likely creates a more favorable environment for viral docking at centromeres, and supports previous findings showing that DNA damage repair at centromeres influences transcription^25,26^.

**Fig. 5.**
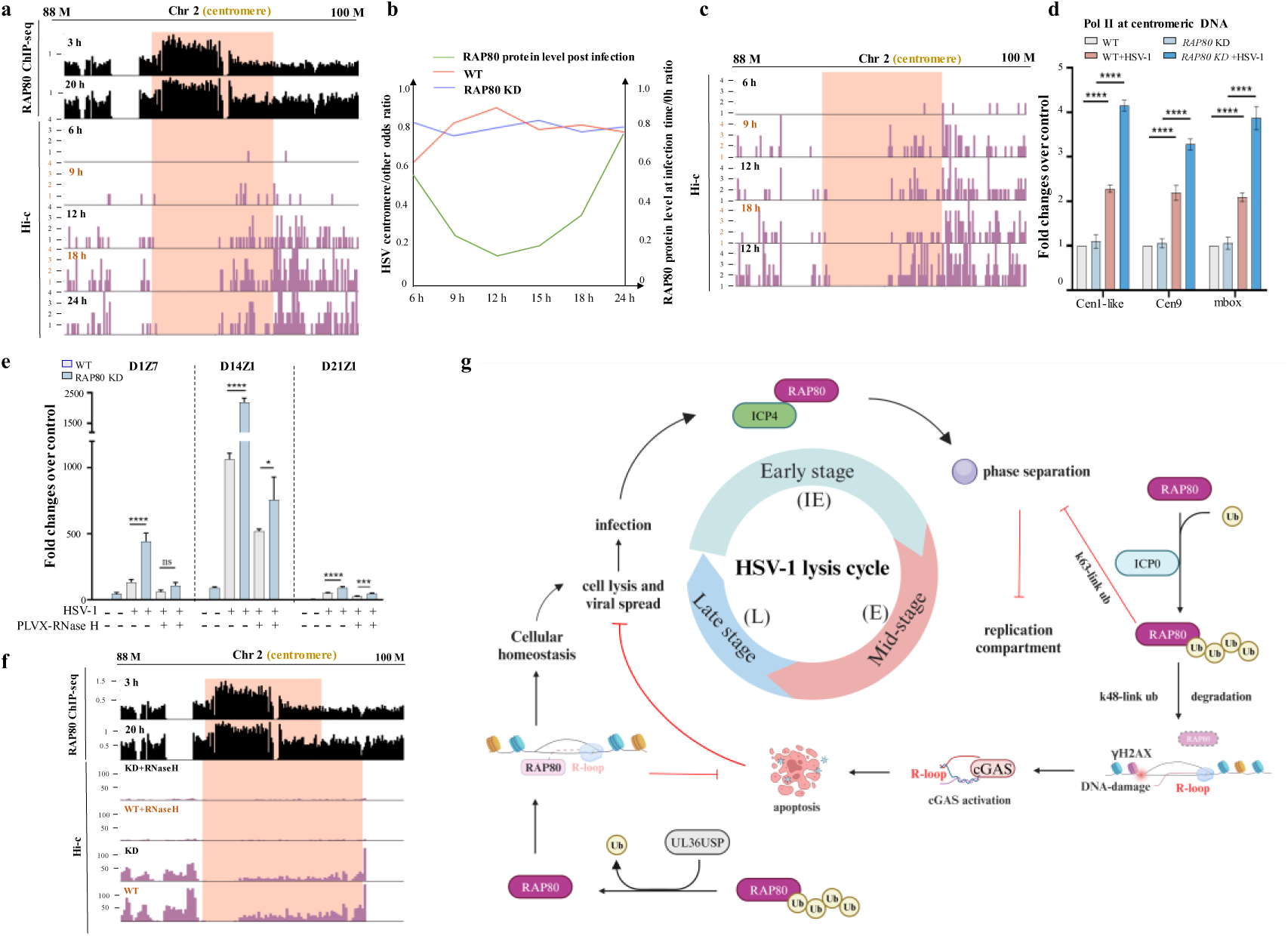
*RAP80* loss promotes viral genome localization to centromeric transcriptionally active regions. **a.** Interactions between HSV-1 and host chromosome 1 (chr.1) near the centromere region. Tracks from the top to the bottom are (1–2) RAP80 enrichment signals in HeLa cells from ChIP-seq data at 3 h (1) and 20 h (2) post-infection. (5–6) Contacts between each 50 kb genomic bin and the HSV-1 genome in WT-HeLa cells are shown at 6 h, 9 h, 12 h, 18 h, and 24 h post-infection. Contacts without mapping quality filters are used. The centromere region is salmon-colored. **b.** The odds ratio for interaction enrichment between HSV-1 and the host genome in the centromere region. An odds ratio of 1 indicates no HSV-1 preference for the centromere region when compared to other regions. A ratio > 1 indicates enrichment in the centromere region, while a ratio < 1 indicates a reduction in that region. Data measured at 6 h, 9 h, 12 h, 15 h, 18 h, and 24 h post-infection are presented. **c.** Similar to **a**, but Hi-C tracks display data from *RAP80* knockdown (KD) HeLa cells. **d.** CUT-tag-qPCR on RNA polymerase II was performed at different centromeric regions in HeLa cells infected with HSV-1 for 20 h (MOI=1). **e.** *RAP80* KD and WT HeLa cells, as well as HeLa cells stably overexpressing RNase H, were infected with HSV-1. Samples were harvested at 20 h post-infection to analyze cenRNA transcripts using RT-qPCR. Data in panels **d, e** are the mean ± standard error of the mean from three independent experiments. *P < 0.05, **P < 0.01 and ***P < 0.001. Ordinary one-way and two-way analysis of variance (ANOVA) and Dunnett’s multiple comparisons tests. Exact P values are provided in source data. **f.** Similar to **a**, but Hi-C track data come from the following 24 h post-infection cells: *RAP80* KD HeLa cells overexpressing RNase H, WT HeLa cells overexpressing RNase, *RAP80* KD HeLa cells, and WT HeLa cells. **g.** The model highlights multifaceted roles of RAP80 in regulating the process of Herpes simplex virus 1 life cycle. RAP80 inhibits HSV-1 transcription in early post-invasion stages by interacting with the viral genome to hinder ICP4 binding to transcription factors. This interaction, mediated by phase separation influenced by multivalent interactions between RAP80’s UIM domain and ICP0/ICP4, is crucial for transcription inhibition. As infection progresses to mid-stages, phase separation dissipates. ICP0 targets RAP80 for proteasomal degradation, disrupting the RAP80-ICP0/ICP4 interaction, dissolving phase separation and facilitating the formation of virus replication compartments, which are essential for viral propagation. In late infection stages, a pivotal change occurs as RAP80 is deubiquitinated by UL36USP, allowing it to play critical roles preserving cellular homeostasis and promoting HSV-1 survival by modulating the R-loop-cGAS-apoptosis axis. This comprehensive analysis of RAP80’s multifaceted roles underscores the dynamic interplay between viral and host factors during HSV-1 infection

RNase H treatment, which suppressed centromeric transcription and RNA polymerase II binding (Fig. 5e, Extended Data Fig.9b), marked decreased virus-host genome interactions at centromeres (Fig. 5f). However, there was little change in interactions at other genome locations (Extended Data Fig.9c). Therefore, RAP80 regulates HSV-1 docking in the host genome by modulating centromere stability and its transcriptional state.

## Discussion

In this study, we delineate complex and stage-specific RAP80 functions during HSV-1 infection, showing dual RAP80 roles throughout viral invasion processes (Fig. 5g). Viral infection involves intricate interactions with host cells. For instance, HSV-1 manipulates host DDR pathways to facilitate its replication. In early post-invasion stages, viral protein ICP4 attract DDR proteins, forming foci structures, but the underlying mechanisms remain poorly understood. We observed that RAP80 interacts with the viral genome, triggering phase separation. In eukaryotic transcription, excessive interactions between transcription factors and mediators may form LLPS compartments, sequestering transcription factors and repressing transcription^29^. Here we observed that RAP80 inhibits early viral transcription by inducing phase separation which competes with ICP4 for binding to its target genes. At middle infection stages, ICP4’s phase separation dissolves into larger mature replication compartments, essential for viral DNA replication. In this study, we investigated regulatory transition dynamics from foci to viral replication compartments and found that ICP0 targets RAP80 for ubiquitination, disrupting RAP80-ICP0/ICP4 interactions and leading to the dissolution of phase separation and the formation of mature replication compartments, highlighting the dynamic interplay between host and viral proteins in the viral reproductive cycle.

During mid-infection stages, ICP0 attaches K48 ubiquitin chains to RAP80 for degradation. However, in late infection stages, UL36USP, which is cleaved from UL36 at approximately 12 h post-infection^22^, deubiquitinates RAP80 and protects it from ICP0-mediated ubiquitination, restoring RAP80 stability and functionality. This previously unknown dynamic regulation step is crucial for late-stage genomic and cellular stability, favoring HSV-1 survival. We also showed that RAP80 is essential for centromere function and R-loop regulation. Failure to properly restore RAP80 causes centromeric dysfunction and R-loop translocation to the cytoplasm, triggering cGAS innate immune responses and promoting apoptosis. Therefore, virus must dynamically adapt and respond to this unstable equilibrium. We highlight the significance of cellular homeostasis in virus-host coevolution by uncovering HSV-1’s dynamic manipulation via RAP80 to counter host defenses.

Moreover, we elucidate pivotal RAP80 roles and mechanisms in preventing R-loop accumulation, thus safeguarding centromere integrity. We found that RAP80 recruits BRCA1 and nucleases to clear R-loops, which is essential for protecting centromeres from DNA damage. RAP80 influences repair pathway choice between RAD51 and RAD52. RAP80 directs RAD51 to DSBs, promoting controlled homologous recombination and genomic stability. To repair DSB, heterochromatin decondensation is required for efficient DSB repair^30^. We showed that HSV-1 infection and *RAP80* KD may prompt heterochromatin remodeling to form more compact chromatin. This is coupled with the observation that HSV-1 infection and *RAP80* deficiency increase RNA polymerase II recruitment. Heterochromatin decondensation facilitates HSV-1 transcription and replication, inducing HSV-1 genome docking at centromeres and impacting virus-host genomic interactions.

Collectively, this study shows that RAP80 is a key non-traditional mediator in virus-host interactions. these findings show the complex coevolution dynamics between viruses and hosts, highlighting essential RAP80 roles in the HSV-1 life cycle, and critically, opening up new avenues for targeting HSV-1 as an oncolytic agent.

## Methods

### Cell culture and viruses

Human embryonic kidney (HEK293T) cells and human cervical cancer cells (HeLa) and BHK cells were from the American Type Culture Collection (ATCC). All cells were cultured in Dulbecco’s Modified Eagle Medium (DMEM) plus 10% fetal bovine serum (Gemini) at 37°C in 5% CO2. The identities of all cell lines were verified by PCR to confirm the absence of mycoplasma contamination.

To generate KD cells, CRISPR-Cas9 gene editing was used. For CRISPR/Cas9-mediated *RAP80* KD in HEK293T and HeLa cells, the following sgRNAs were used:

SgRAP80 #1-F: CACCGGTCGAATAGAGCAAAGTGTT; SgRAP80 #1-R: AAACAACACTTTGCTCTATTCGAC; SgRAP80 #2-F: CACCGTAGCGGAAGCATCAGAAGGC; SgRAP80 #2-R: AAACGCCTTCTGATGCTTCCGCTA; SgRAP80 #3-F: CACCGATTGTGATATCCGATAGTGA; SgRAP80 #3-R: AAACTCACTATCGGATATCACAAT;

Study viruses included the wild-type (WT) HSV-1 strain 17 and a mutant virus *ΔICP0-HSV-1* (ICP0 enzyme activity deletion mutant, strain 17). Viruses were amplified and titrated in BHK cells. Cells were infected for 1 h in fresh media, and then the inoculum was replaced with DMEM plus 2% fetal bovine serum.

### Antibodies

The following antibodies were used: Anti-Flag (M2, F3165, Sigma), anti-HA (HA-7, H9658, Sigma), anti-Myc (M047-3, MBL), anti-His (D291-3x, L), anti-β-actin (PM053, Abclonal), anti-RAP80 (A7244, Abclonal), anti-ICP0 (sc-53070, Santa Cruz), anti-ICP4 (sc-69809, Santa Cruz), anti-ICP8 (sc-53329, Santa Cruz), anti-RNA polymerase II S5P (ab5408, Abcam), anti-cGAS (sc-515777, Santa Cruz), and anti-CENP-A (A15995, Abclonal).

### Immunofluorescence

Cells at an appropriate density (2-3×10^6^) were inoculated into 6-well culture plates plus sterile cover slips. After culture or transfection for an appropriate time, cover slips with adhered cells were washed three times in precooled phosphate buffered saline (PBS), fixed in 4% paraformaldehyde for 15 min, and cells perforated with 1% Triton X-100 for 5 min. Cover slips were washed three times in precooled PBS and then blocked in 1% bovine serum albumin (BSA, NEB B9000S) at room temperature for 1 h. A primary antibody (1:100 diluted in BSA solution) was then added and incubated at room temperature for 1 h or 4°C overnight. Cover slips were then washed three times in precooled PBS to remove non-binding antibodies. Then, a FITC- or TRITC-labeled secondary antibody (1:100 diluted in BSA) was added and cover slips incubated at room temperature in the dark for 1 h. Slips were then washed three times in precooled PBS to remove excess secondary antibody. Next, sealing agent containing 4′,6-diamidino-2-phenylindole (DAPI) was dropped onto slides, the edges sealed with nail polish, and fixed. Slides were temporarily stored at 4°C away from light, after which image were observed and analyzed using a Zeiss LSM 710 laser confocal microscope.

### The Hi-C protocol

Cells were fixed in 1.75% formaldehyde, quenched in a 0.1% BSA solution, washed, and resuspended in 200 μL ice-cold wash buffer (10 mM Tris (pH 8.0), 10 mM NaCl, and 0.1 mg/mL BSA with 20 μL of a protease inhibitor (Sigma P8340). Then, cells were treated in sodium dodecyl sulfate (SDS) and Triton X-100 and digested with the restriction enzyme *MboI*. After deactivating the enzyme, cells underwent biotinylation and ligation reactions, after which 50 ng of the Hi-C sample was used for library preparation. The final library was sequenced on a NovaSeq 6000 (Illumina) platform according to manufacturer’s instructions. Paired-end 150 bp reads were generated.

### Hi-C data processing

To simultaneously consider both host and HSV genomes, we created a fusion reference genome by combining GRCh38 and NC_001806.2 sequences. We then processed Hi-C data using the distiller-nf pipeline (https://github.com/open2c/distiller-nf). Briefly, we mapped Hi-C reads to the fusion reference genome using BWA (v.0.7.17). Next, we used Pairtools (v.0.2) to filter out reads with MAPQ scores < 30 to remove PCR duplications and extract Hi-C contacts. To show interactions between HSV-1 and the host genome, we divided the host genome into uniform bins of 500 kb in size and calculated the number of contacts between each bin and any HSV region. To visually represent interactions between HSV, host centromeres, and surrounding areas, we created plots using contact matrices generated without MAPQ filtering. We also used these matrices to quantitatively assess HSV-1 enrichment in centromeric regions. Specifically, we calculated HSV-1-related contact ratios in centromeric regions to those in remaining regions, resulting in an odds ratio to measure enrichment.

### Immunoprecipitation and immunoblotting

Cells were collected at 4°C for 5 min at 3,000 rpm and washed once in pre-cooled PBS. Cells were then transferred to a tube and 1 mL of pre-cooled lysis buffer and a protease inhibitor (final concentration=20 µM) added and tubes rotated at 4°C for 1 h. Lysates were then sonicated at 200 W for 5 s each over five intervals; 10 times in total. Lysates were then centrifuged at 13,000 rpm for 15 min at 4°C to remove cellular debris, after which 30 µL of supernatant was taken. Then, 1 µg of corresponding or homologous IgG (control) antibody was added to the supernatant and incubated at 4°C for 3–4 h. Add 30 µL 50% Protein G agarose column material, incubate at 4°C for 3-4 h, and enrich the immunoprecipitation complex. This was left on ice for 3 min, re-centrifuged at 4°C for 5 min at 3,000 rpm, the supernatant discarded, and the pellet washed three times in precooled lysis buffer to remove non-specific proteins binding to the antibody. Tubes were then centrifuged at 4°C for 5 min at 3,000 rpm, supernatants discarded and 30 µL of 2×SDS loading buffer added, and then incubated at 96°C for 10 min and stored at -20°C.

For immunoblotting, proteins were fractionated using SDS-polyacrylamide gel electrophoresis, followed by transfer to nitrocellulose filter membranes (GE Healthcare, USA). Once protein transfer was complete, membranes were blocked in 5% milk, then sequentially incubated with primary and secondary antibodies at appropriate dilutions. Protein signals were detected and quantified using the Odyssey Infrared Imaging System (LI-COR Biosciences, USA) or the AI 600 Ultra-Sensitive Multifunction Imager (GE, USA).

### Protein purification and *in vitro* phase separation

A target plasmid was transformed into *Escherichia coli* BL21 competent cells, which were grown until a specific optical density was reached, after which isopropyl β-D-1-thiogalactopyranoside (IPTG) (final concentration=0.5 mM) was added to induce protein expression, and cells finally cultured at 18° for 16 h to 20 h at 220 rpm. Proteins were purified on an AKTA FPLC protein purification system. To form dorplots *in vitro*, purified proteins were added to *in vitro* phase separation buffer (50–150 mM salt and 20%–25% PEG8000) and observed under an inverted microscope.

### Quantitative PCR with Reverse Transcription (qRT-PCR)

Total RNA was extracted in Trizol (Invitrogen), after which cDNAs were synthesized from 2 µg total RNA with random or oligo (dT) primers using FastQuant RT Kits (TIANGEN). Quantitative PCR was performed using SYBR Green Master Mix (YEASEN) in a LightCycler 480 instrument (Roche) and analyzed in LightCycler 480 software using the 2^-ΔΔCt^ method.

### CUT-tag assays

Approximately 500,000 cells were washed twice in 100 µL wash buffer and then mixed 90 µL cells with 10 µL activated NovoNGS ConA Beads. A primary antibody was diluted (1:50–1:100) in antibody buffer, add 50 µL of pre-cooled primary antibody diluent, and incubate overnight at 4°C. Magnetic beads were separated from the liquid and the supernatant carefully aspirated. Then, 100 µL Dig wash buffer (1:100 ratio) was added for secondary antibody dilution, resuspend all ConA magnetic beads-cells, gently mix and blow, and incubate at room temperature for 1 h. After washing twice in Dig-wash buffer, cells were incubated with 1 µL of pAG-Tn5 for 1 h. After resuspending cells, 300 µL of Tagmentation Buffer was added and gently blow the resuspended cells, and incubated at 37°C for 1 h. Finally, the chromatin complex was digested in a solution containing 10 µL of 0.5 M EDTA, 3 µL of 10% SDS, and 2.5 µL of 20 mg/mL Proteinase K at 55°C for 1 h. Transposed DNA fragments were purified using a DNA extraction solution (phenol: chloroform: isoamyl alcohol=25:24:1).

### Metaphase spreads

Cells were harvested by trypsinization, supernatants completely removed and resuspended in 5 mL of 0.075 M KCl. Cells were incubated for up to 30 min at 37°C, inverting tubes to keep cells suspended. Dropwise, 500 µL of fixative was added while cells were slowly and gently mixing. When ready, cells were centrifuged (1,000 rpm), 9 mL of fixative was removed and cells resuspended in the remaining 1 mL (may vary depending on cell number). Resuspended cells were dropped (from a couple of inches) onto the ends of a wet slide, the nuclei washed in fresh fixative, and slides placed for 1 min on a humidified 70°C–80°C heating block. Finally, DAPI was dropped onto slides and the edges sealed with nail polish and fixed.

### Neutral comet assays

Cells at 1 x 10^5^/mL (in 10 µL) were combined with molten LMAgarose (100 µL at 37°C) at a 1:10 (v/v) ratio, and immediately 70 µL was pipetted onto CometSlides^TM^. The side of the pipette tip was used to spread agarose/cells over the sample area. Slides were placed flat at 4°C in the dark for 10–30 min. A 0.5 mm clear ring appears at edge of CometSlide^TM^ area. Increasing gelling time to 30 min improves cell adherence in high humidity environments. For added sensitivity, slides were immersed in 4°C lysis solution for 1 h or overnight. After removal from lysis buffer, excess buffer was drained and slides were gently immersed in 50 mL of 4°C 1× neutral electrophoresis buffer for 30 mins. In the CometAssay® ES unit, 850 mL of 4°C 1× neutral electrophoresis buffer was added, the slides placed in the electrophoresis slide tray, and covered with a slide overlay tray. The power supply was set to 21 volts and applied for 45 min at 4°C. Excess neutral electrophoresis buffer was drained and slides gently immersed in DNA precipitation solution for 30 min at room temperature. Slides were then immersed in 70% ethanol for 30 min at room temperature, after which they were dried at 37°C for 10–15 min. Drying brings cells into a single plane to facilitate observations. Samples may be stored at room temperature with a desiccant prior to scoring. 5µg/mL PI, RT, dark, staining 10 min. ddH_2_O rinse and photograph.

### Statistical analyses and reproducibility

Data were analysed for statistically significant differences using GraphPad Prism 8. Gene expression values were log2-transformed. Statistical tests are indicated in the Fig. legends. ata are presented as mean + s.e.m. of three independent experiments. **P < 0.01; ***P < 0.001; The data analysis was conducted using the Ordinary one-way and two-way analysis of variance (ANOVA) and Dunnett’s multiple comparisons test. Exact P values are provided in the Source Data. Scale bars, 5 μm. nd stimulus. Normality and sphericity of data were assumed and not tested. *P < 0.05, **P < 0.01, ***P < 0.001.

## Resource availability

### Lead contact

Further information and requests for resources and reagents should be directed to and will be fulfilled by the lead contact, Xiaofeng Zheng.

## Data and code availability

The ChIP-seq data have been deposited to the GEO Dataset: https://www.ncbi.nlm.nih.gov/geo/query/acc.cgi?acc=GSE281836

## Acknowledgements

We sincerely thank Prof. Zhengfan Jiang from Peking University for providing BHK cells, Prof. Chunfu Zheng from Fujian Medical University for providing HSV-1 strain 17 and a mutant virus ΔICP0-HSV-1, and T. Li help with bioinformatics analysis. We thank the National Center for Protein Sciences and the Core Facilities of Life Sciences at Peking University, particularly hongxia Lv, Yinghua Guo, Qi Zhang, Wenling Gao, Siying Qin, Liying Du, Huan Yang, Jia Luo, Feifei Cheng, and Guilan Li for technical help.

## Author contributions

J. Wang and X. Zheng designed this study. J. Wang conducted the experiments. J. Wang analyzed the data and wrote the manuscript. Y. Deng and Y. Chi performed HiC related experiments and analyzed the data. D. Zhu helped with experiments of cell culture, knockout cell line construction, and viral amplification. X. Zheng and D. Xing supervised this study and wrote the manuscript.

