## Supplementary material for "HSV-1 orchestrates host RAP80 ubiquitination by ICP0 and UL36USP to promote viral survival": Supplemenntary Figures 1-9

### Extended data

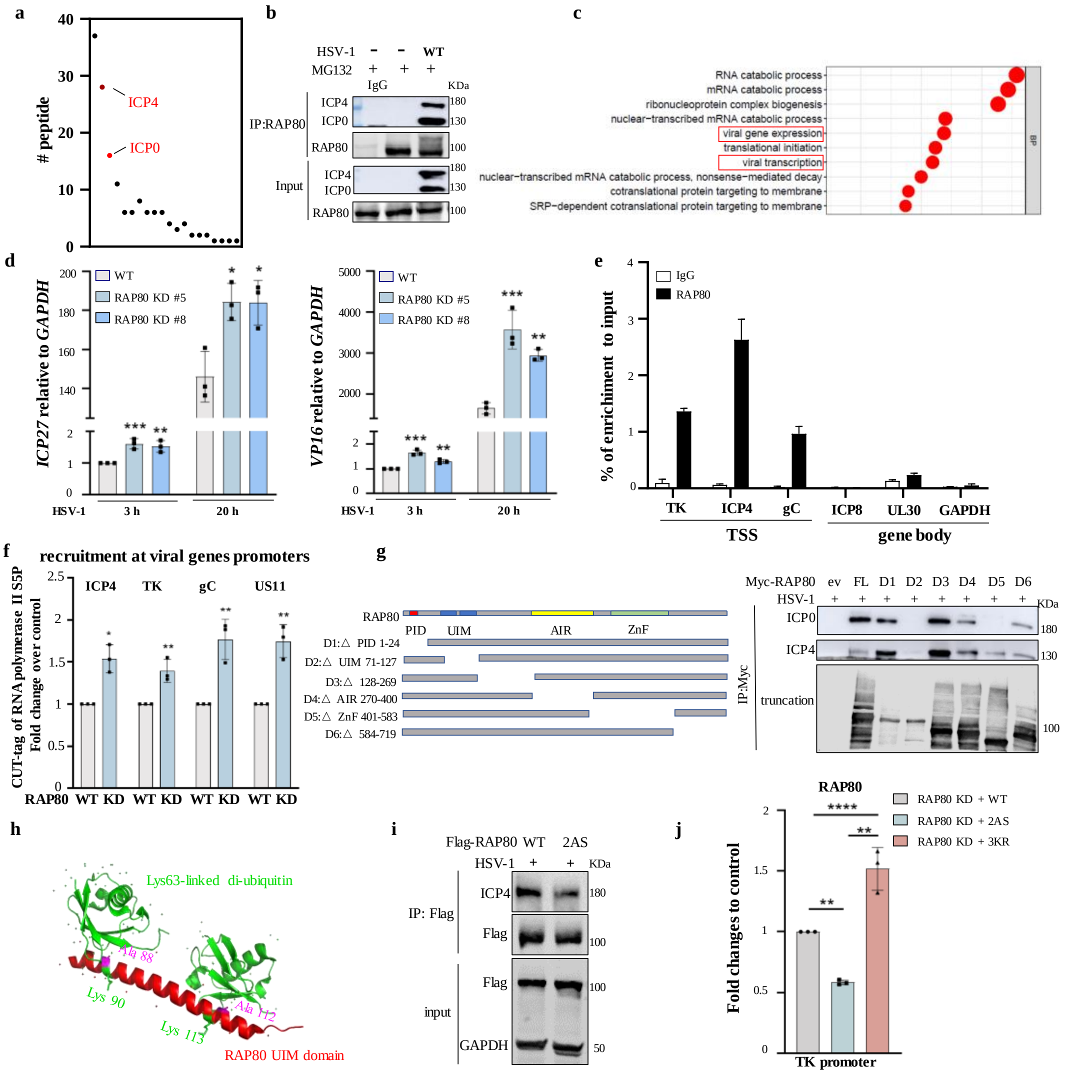


**Extended Data Fig.1 RAP80 interacts with ICP4 and inhibits HSV-1 transcription.**

**a**. IP-MS assay showing specific RAP80-associated viral proteins. HEK293T cells harboring Flag-RAP80 were infected with HSV-1 (MOI=1) for 3 h, and then IP was performed with anti-Flag M2-agarose beads. Precipitated proteins were analyzed by mass spectrometry.

**b**. HeLa cells were infected with HSV-1 (MOI=1) for 3 h, and IP performed with an anti-RAP80 antibody, after which viral protein ICP4 and ICP0 associations with RAP80 were assessed by western blotting using anti-ICP0 and anti-ICP4 antibodies.

**c.** To elucidate the RAP80 interaction landscape with the HSV-1 genome, HeLa cells were infected with HSV-1 (MOI=1). Following a 3 h incubation, cells were subjected to ChIP-seq analysis using an RAP80-specific antibody to obtain ChIP DNA. Peaks, representing RAP80 binding sites, were mapped and visualized in the HSV-1 genome.

**d**. *RAP80* knockdown (KD) effects on *ICP27* and *VP16* expression at early (3 h) and late (20 h) viral infection stages were investigated using qRT-PCR assays in HeLa cells.

**e**. RAP80 binding to promoters and gene bodies in viral genes was assessed in HeLa cells using ChIP assays and a RAP80 antibody.

**f**. *RAP80* KD effects on RNA polymerase II binding to viral gene promoters were investigated in

WT and *RAP80* KD HeLa cells. After infection with HSV-1 (MOI=1) for 20 h, CUT-tag assays were performed with the RNA polymerase II S5P antibody to analyze promoter occupancy for the viral genes *ICP4, TK, gC,* and *US11*.

**g**. Co-IP assays were conducted to identify critical domain(s) in RAP80 responsible for associations with the viral proteins ICP0 and ICP4. HEK293T cells expressing Myc-tagged RAP80 and its truncated mutants were infected with HSV-1 (MOI=1) for 3 h, and then co-IP assays performed with indicated antibodies.

**h**. The three-dimensional structure of the RAP80 UIM domain in complex with Lys63-linked di-ubiquitin was predicted using Alphafold2. A88 and A113 residues in RAP80 are highly conserved among UIMs and are important for function. The RAP80 UIM domain contains two lysine residues, K90 and K112, which are potential ubiquitination sites.

**i**. HEK293T cells expressing Flag-tagged RAP80 and the *2AS* mutant were infected with HSV-1 (MOI=1) for 3 h, after which IP with an anti-Flag antibody was performed to assess ubiquitin binding effects, as mediated by the UIM domain, to viral protein ICP4.

**j**. The comparative analysis of RAP80 recruitment to viral gene promoters was assessed in HeLa cells for WT RAP80 and its mutants (*2AS* or *3KR*) using ChIP assays with an antibody specific to RAP80.

Data in panels **d, f**, and **j** are the mean ± standard error of the mean from three independent experiments. *P < 0.05, **P < 0.01, and ***P < 0.001. Ordinary one-way and two-way analysis of variance (ANOVA) and Dunnett's multiple comparisons tests. Exact P values are provided in source data. Scale bars, 5 μm.

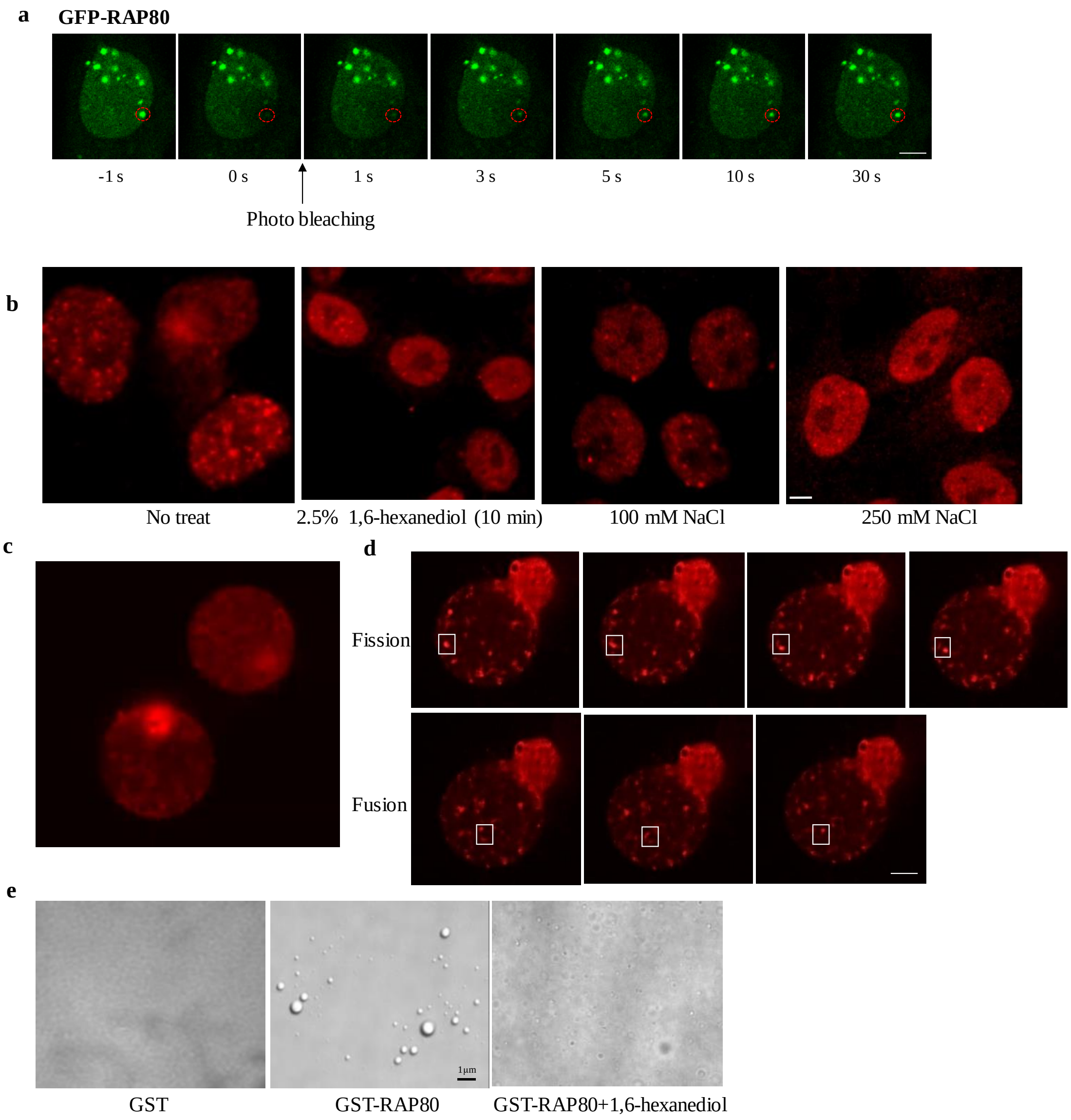


**Extended Data Fig.2 RAP80 undergoes liquid–liquid phase separation.**

**a**. GFP-RAP80 dynamics were examined by photo-bleaching analysis coupled to live-cell imaging studies in HeLa cells expressing GFP-tagged RAP80. Sequential live-cell images were captured at precise intervals following the initiation of photo-bleaching (time zero).

**b**. Immunofluorescence images showing the punctate localization of endogenous RAP80 in HeLa cells treated with different NaCl and 1,6-hexanediol concentrations.

**c, d.** Live-cell video microscopy was used to document dynamic fission and fusion processes of mCherry-tagged RAP80 puncta in living cells.

**e**. Confocal microscopy showing LLPS droplet formation of purified GST-tagged RAP80 in *in vitro* phase separation buffer or in the presence of 1,6-hexanediol (Scale bar, 1 μm).

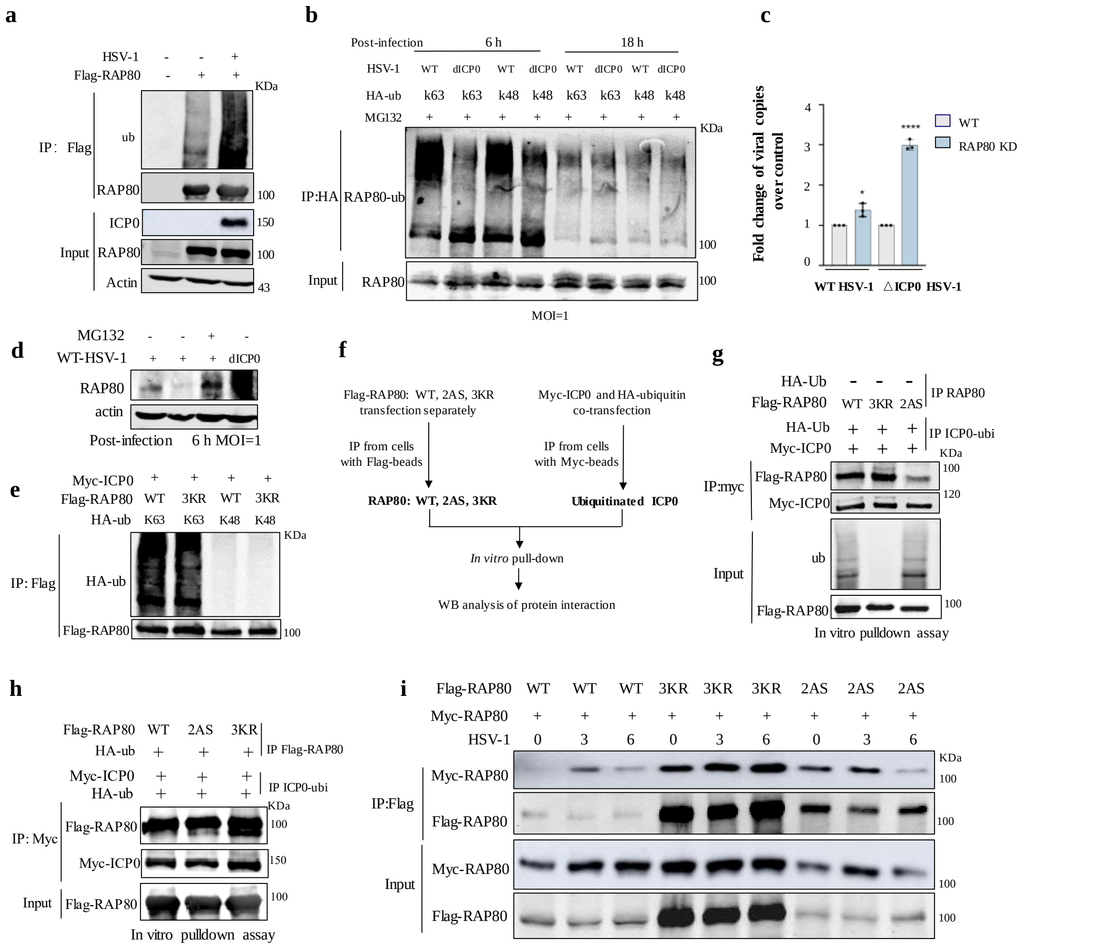


**Extended Data Fig.3 ICP0 disrupts RAP80 multivalent interactions by ubiquitinating lysine residues in the RAP80 UIM domain.**

**a**. HEK293T cells harboring Flag-tagged RAP80 were infected with HSV-1 (MOI=1) for 6 h, after which IP was performed with an anti-Flag M2 antibody. Immunoprecipitates were analyzed by western blotting with indicated antibodies.

**b**. HSV-1 and ICP0 activity-deficient HSV-1 (*ΔICP0 HSV-1*) effects on the K63- or K48-linked polyubiquitination of RAP80 were examined in HEK293T cells. Cells transfected with indicated plasmids were infected with HSV-1 or *ΔICP0 HSV-1* and lysed at 6 h and 18 h post-infection under denaturing conditions. RAP80 ubiquitination levels were examined using IP assays with indicated antibodies.

**c**. Viral cDNAs were synthesized and measured by q-RT-PCR in WT and *RAP80* KD HeLa cells infected with WT HSV-1 and *ΔICP0 HSV*-1 (MOI=1) for 6 h. Data are presented as the mean ± standard error of the mean from three independent experiments. *P < 0.05 and ****P < 0.0001. Data analysis was conducted using ordinary two-way analysis of variance (ANOVA) and Dunnett's multiple comparisons tests. Exact P values are provided in source data.

**d**. HeLa cells treated with MG132 were infected with WT HSV-1 and *ΔICP0 HSV-1* (MOI=1), and at 6 h post-infection, endogenous RAP80 levels were analyzed by immunoblotting.

**e**. ICP0 effects on the K63- and K48- linked ubiquitination of RAP80 were detected in HEK293T cells. Cells transfected with indicated plasmids were infected with HSV-1 for 6 h and subjected to IP assays using specific antibodies.

**f**. Diagram showing *in vitro* pulldown assays in **g**, **h**.

**g**. UIM domain effects on the ICP0-RAP80 multivalent interaction were examined using *in vitro* pulldown assays. WT, mutant *RAP80*, and ubiquitinated ICP0 proteins were precipitated from HEK293T cells following the procedure in (f) and subjected to *in vitro* pulldown assays to examine interactions between ICP0 and RAP80.

**h**. Ubiquitination effects at 3K sites (K75, 90, and 112) on the multivalent interaction between ICP0 and RAP80 were examined using *in vitro* pulldown assays following the procedure described in (f).

**i**. The impact of HSV-1 on the interaction between WT RAP80 and mutant proteins was investigated in HEK293T cells co-transfected with indicated plasmids. Following infection with HSV-1 (MOI=1), cells were harvested at 0 h, 3 h, and 6 h post-infection, and subjected to co-IP assays using indicated antibodies.

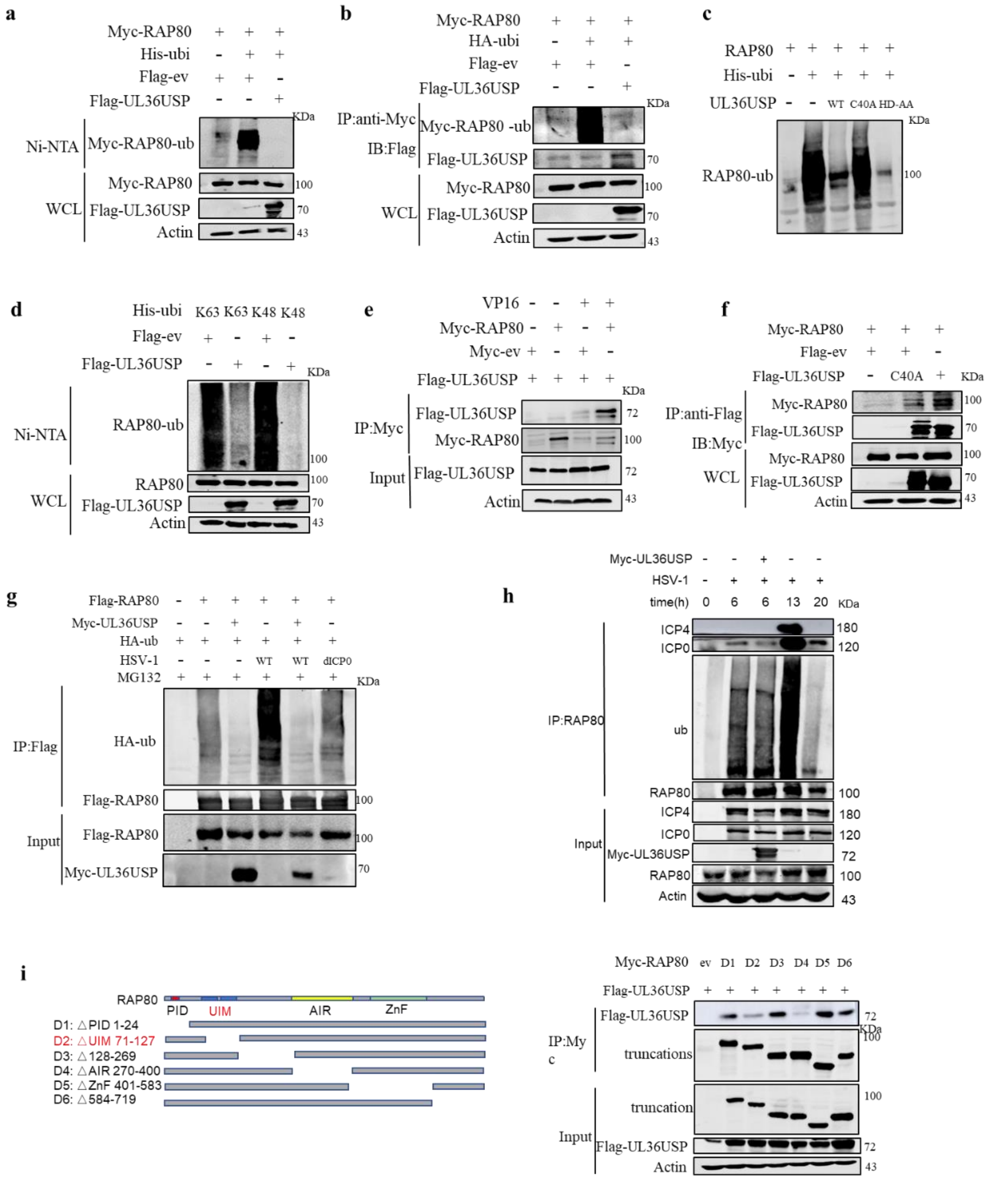


**Extended Data Fig.4 UL36USP deubiquitinates RAP80 and blocks its interaction with ICP0.**

**a**, **b**. The deubiquitination effects of UL36USP on RAP80 were examined in HEK293T cells. Cells transfected with indicated plasmids were subjected to His-ubiquitin pulldown (a) or denaturing IP (b) assays with indicated antibodies.

**c**. UL36USP deubiquitinase effects on RAP80 ubiquitination were assessed in HEK293T cells expressing WT or UL36USP enzyme-deficient *C40A* and *HD-AA* (the two other catalytic residues in the enzyme activity center were mutated) mutants using His-ubiquitin pulldown assays.

**d**. UL36USP effects on the K48- and K63- ubiquitination of RAP80 were examined by His-ubiquitin pulldown assays in HEK293T cells transfected with specified plasmids.

**e**. DNA damage effects on the interaction between UL36USP and RAP80 were assessed using Co-IP assays in HEK293T cells transfected with indicated plasmids.

**f**. Co-IP assays were performed to examine the interaction between RAP80 and WT UL36USP or the enzyme-deficient mutant *C40A* in HEK293T cells transfected with indicated plasmids.

**g**. UL36USP effects on RAP80 ubiquitination in HEK293T cells infected with HSV-1 or *ΔICP0 HSV-1* (MOI=1). Cells pre-treated with MG132 were harvested at 6 h post infection and subjected to denaturing IP assays to examine RAP80 ubiquitination.

**h**. Viral infection effects on RAP80 ubiquitination and its interaction with viral proteins ICP0/4 in HEK293T cells at different post-infection stages. Cells transfected with specific plasmids were infected with HSV-1 and harvested at 0 h, 6 h, 13 h, and 20 h post-infection. RAP80 ubiquitination levels and RAP80-ICP0/4 interactions were then assessed.

**i**. Mapping the RAP80 domain responsible for its interaction with UL36USP. HEK293T cells transfected with various Flag-*RAP80* mutation constructs were subjected to co-IP assays.

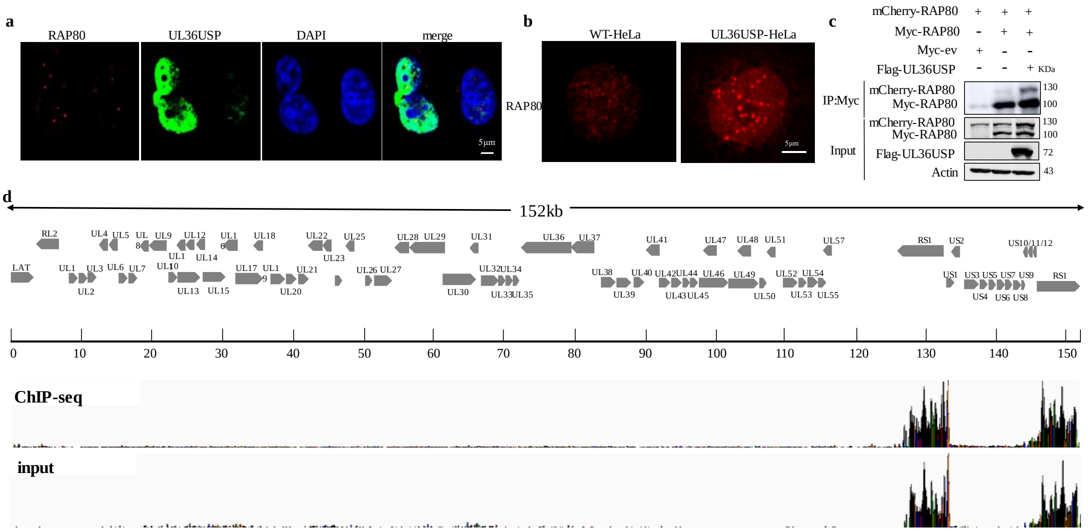


**Extended Data Fig. 5** UL36USP promotes RAP80 function.

**a**, **b**. UL36USP effects on the phase separation of RAP80 were examined in HeLa cells either transiently transfected with UL36USP (a) or stably overexpressing UL36USP (b). RAP80 distribution by immunofluorescence microscopy following immunostaining with a specific anti-RAP80 antibody.

**c**. UL36USP effects on RAP80 self-associations were investigated using co-IP assays in HEK293T cells co-transfected with Myc-RAP80 and mCherry-RAP80.

**d**. RAP80 genomic recruitment sites were mapped using ChIP-sequencing analysis, using RAP80 ChIP DNA obtained from HeLa cells at 20 h post-infection (MOI=1). Peaks, corresponding to RAP80 binding sites in the HSV-1 genome, were graphically represented to illustrate RAP80 spatial distribution and enrichment patterns.

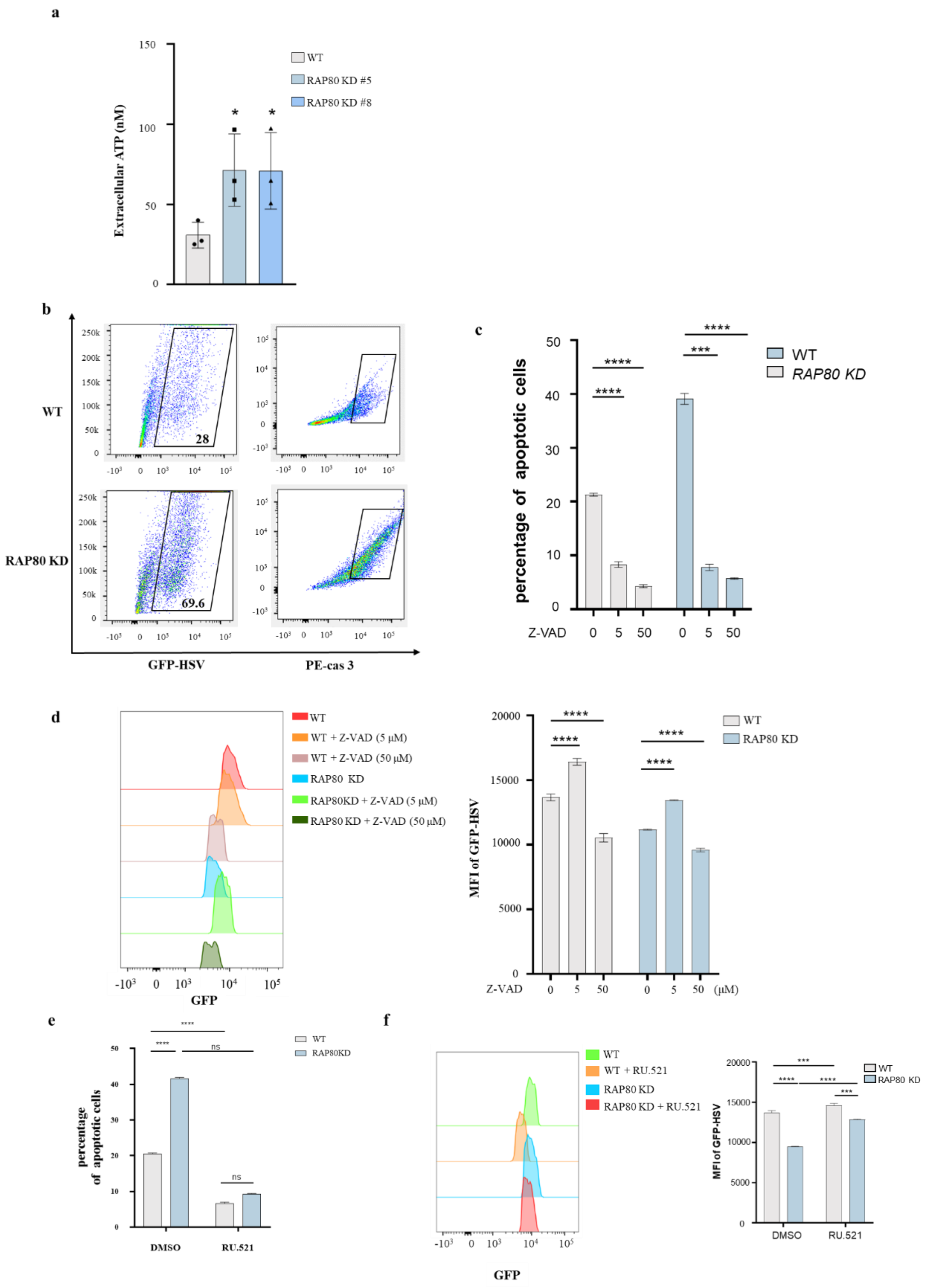


**Extended Data Fig.6 The RAP80-R-loop-cGAS-apoptosis axis serves as a regulatory mechanism controlling viral reproduction.**

**a**. ATP assays were conducted to evaluate danger signal release in conditioned media from HSV1-infected WT or *RAP80*-depleted HeLa cells undergoing cell death.

**b**. *RAP80* knockdown (KD) effects on HSV-1-induced apoptosis were investigated using flow cytometry in HeLa cells at 20 h post-infection with GFP-HSV-1.

**c**. Z-VAD concentration effects on apoptosis in HSV-1-infected HeLa cells were evaluated using flow cytometry. WT and *RAP80*-depleted cells were infected with HSV-1 and subsequently treated with Z-VAD, a pan-caspase inhibitor that inhibits apoptosis, at 0, 5, and 50 μM concentrations. At 20 h post-infection, cells were harvested and subjected to a double-staining protocol using annexin V and propidium iodide (PI) to assess apoptosis.

**d**. The effects of varying Z-VAD concentrations on HSV-1 amplification were investigated. WT and *RAP80*-depleted cells were infected with GFP-tagged HSV-1 and subsequently exposed to 0, 5, and 50 μM Z-VAD concentrations. Mean fluorescence intensity (MFI) was measured using flow cytometry.

**e**. RU.521 effects on apoptosis in HSV-1-infected HeLa cells were evaluated using flow cytometry. WT and *RAP80*-depleted cells were infected with HSV-1 (MOI=1), followed by treatment with the cGAS inhibitor RU.521 (5 μM). At 20 h post-treatment, cells were collected and apoptosis induced by viral infection was evaluated by staining with annexin V and PI.

**f**. RU.521 effects on HSV-1 amplification were examined in HeLa cells as previously described (e). Flow cytometry was used to quantify MFI.

Data in panels **a, c-e** are presented as the mean ± standard error of the mean from three independent experiments. *P < 0.05, **P < 0.01, and ***P < 0.001. Data analysis was conducted using ordinary one-way and two-way analysis of variance (ANOVA) and Dunnett's multiple comparisons tests. Exact P values are provided in source data.

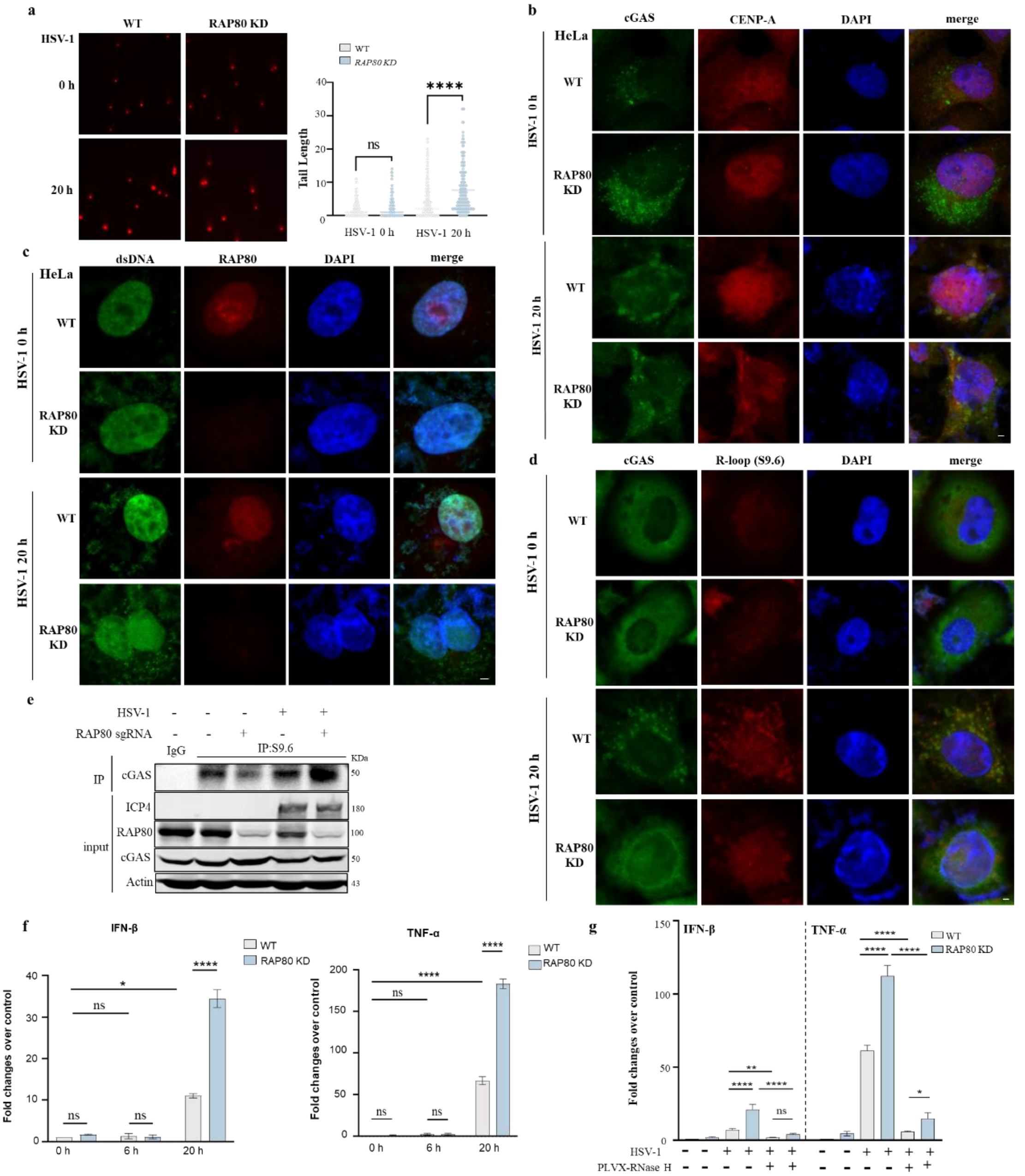


**Extended Data Fig.7 The RAP80-mediated R-loop-cGAS-apoptosis axis has a regulatory role in modulating viral replication.**

**a.** Viral infection effects on RAP80-mediated genome integrity were assessed by neutral comet assays. WT and *RAP80*-depleted HeLa cells were infected with HSV-1 for 20 h and assays performed. Approximately $2\times10^5$ cells were counted/group.

**b–d**. The spatial colocalization of different proteins or dsDNA was examined using immunofluorescence microscopy. WT and *RAP80*-depleted HeLa cells were infected with HSV-1 for 20 h, and IF analyses conducted using specific antibodies to visualize colocalization patterns. The representative images show the spatial relationships between cGAS and CENP-A (b); dsDNA and RAP80 (c); and cGAS with S9.6 (d).

**e**. The interaction between cGAS and R-loops was assessed in HeLa cells following HSV-1 infection. WT and *RAP80*-depleted HeLa cells were infected with HSV-1 (MOI=1), after which cells were harvested at 20 h post-infection and subjected to IP assays using specific antibodies to determine cGAS associations with R-loops.

**f**, **g**. *RAP80* knockdown (KD) effects on activated innate immune responses were examined by RT-qPCR. IFN-β and TNF-α expression levels were measured in WT and *RAP80*-depleted HeLa cells at various stages following HSV-1 infection (**f**) or in HSV-1-infected cells with/without RNase H treatment (**g**).

Data in panels **a, f,** and **g** are presented as the mean ± standard error of the mean from three independent experiments. *P < 0.05, **P < 0.01 and ***P < 0.001. Data analysis was conducted using ordinary one-way and two-way analysis of variance (ANOVA) and Dunnett's multiple comparisons tests. Exact P values are provided in source data. Scale bars, 5 μm.

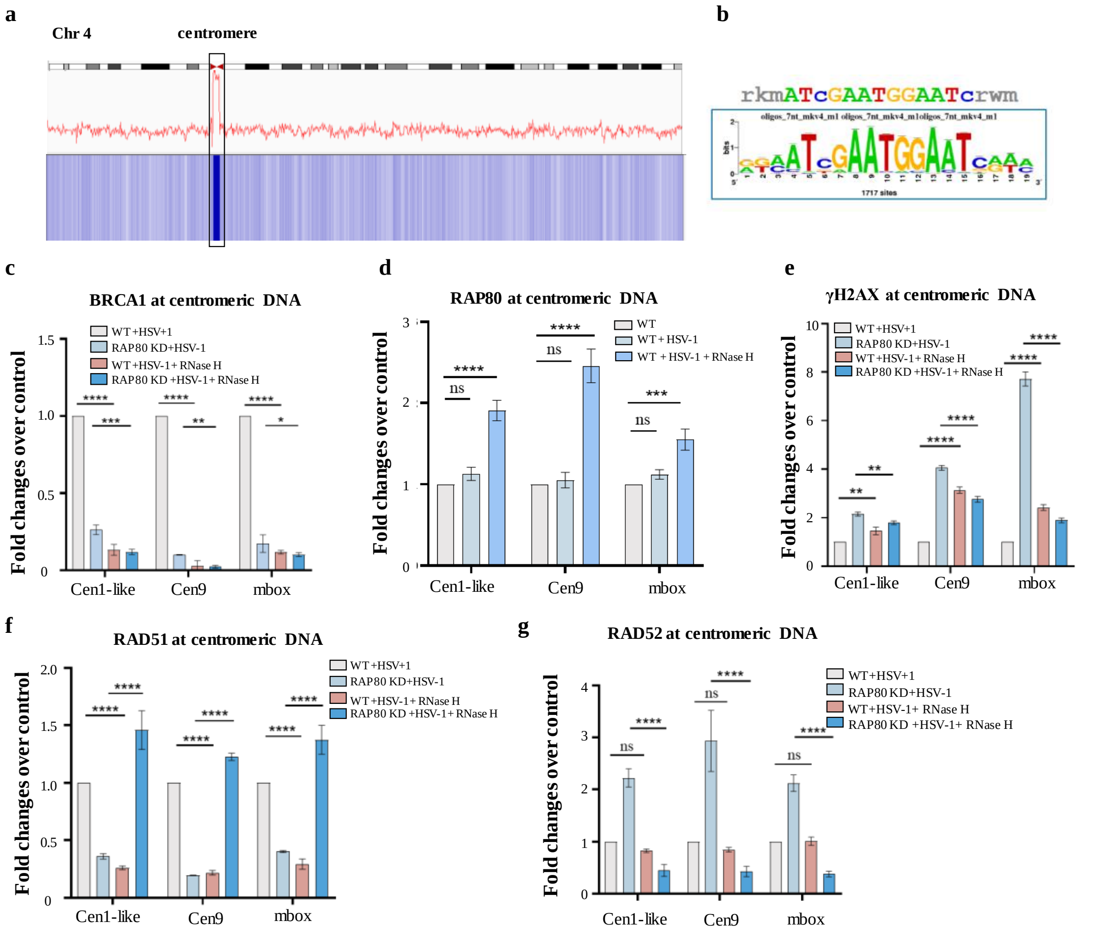


**Extended Data Fig. 8 R-loop accumulation induces centromeric instability in chromosomes.**

**a**. Plot showing the spatial distribution of RAP80 peak locations across the hg38 genome assembly.

**b**. A motif analysis of RAP80 binding sites across the entire HeLa cell genome.

**c–g.** Multiple protein enrichment, including BRCA1 (**c**), RAP80 (**d**), γH2AX (**e**), RAD51 (**f**), RAD52 (**g**) at centromeric DNA regions, was evaluated in WT and *RAP80* KD HeLa cells. CUT-tag-qPCR assays were used to characterize protein binding profiles to specific genes (*Cen1-like, Cen9,* and *mbox*). Experiments were conducted under different conditions, including HSV-1 infections and ribonuclease H (RNase H) treatments.

Data in panels **c-g** are presented as the mean ± standard error of the mean from three independent experiments. *P < 0.05, **P < 0.01 and ***P < 0.001. Data analysis was conducted using ordinary two-way analysis of variance (ANOVA) and Dunnett's multiple comparisons tests. Exact P values are provided in source data.

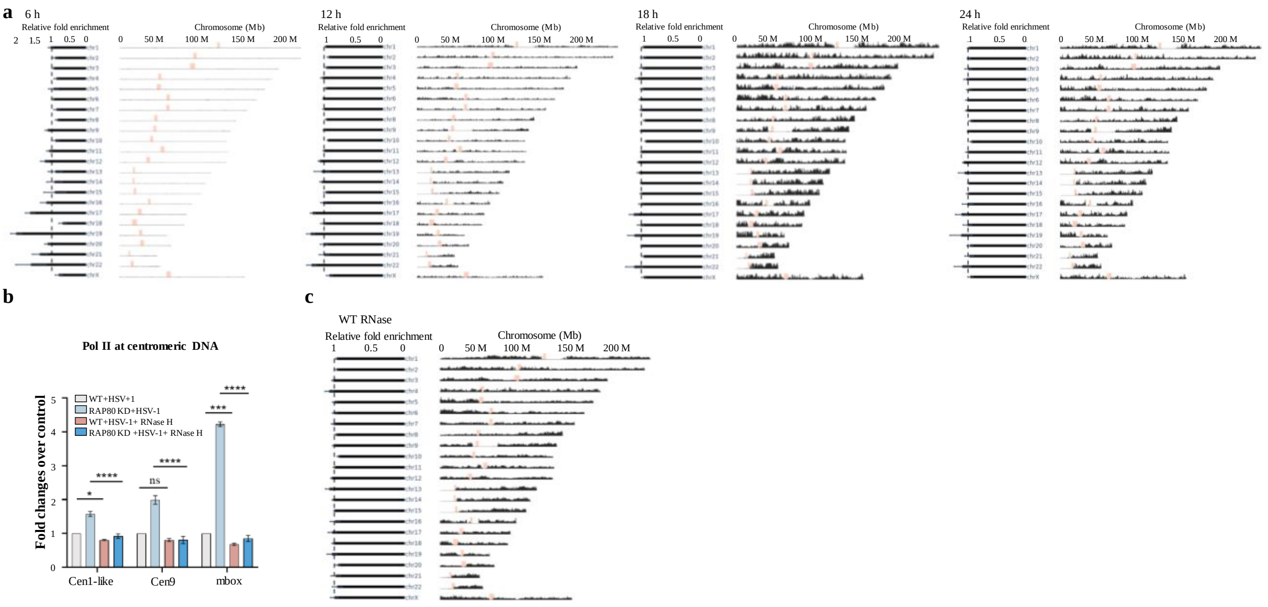


**Extended Data Fig. 9 The distribution of HSV-1 in host chromosome.**

**a.** Interactions between HSV-1 and host chromosomes in HeLa cells at 6 h, 12 h, 18 h, and 24 h post-infection. For each panel, the left horizontal bar displays the ratio of contacts between each chromosome and the HSV-1 genome relative to the intra-chromosomal contacts of the chromosome itself, normalized by the median of all chromosomes. The vertical dashed line represents the median. A value > 1 indicates that this chromosome is enriched in viral interactions relative to other chromosomes, while a value < 1 indicates a reduced interaction. The right side shows the contacts between each 500 kb host genomic region and the viral genome, normalized by the total number of contacts for the chromosome that contains this genomic region.

**b.** CUT-tag-qPCR on RNA polymerase II was performed at different centromeric regions in HeLa cells infected with HSV-1 for 20 h (MOI=1) with overexpressing RNase H. Data are presented as the mean ± standard error of the mean from three independent experiments. *P < 0.05 and ***P < 0.001. Data analysis was conducted using ordinary two-way analysis of variance (ANOVA) and Dunnett's multiple comparisons tests. Exact P values are provided in source data.

**c**. Similar to panel **a**, but shows 24 h post-infection results from WT HeLa cells overexpressing RNase H.
